# PRDM16 functions as a co-repressor in the BMP pathway to suppress neural stem cell proliferation

**DOI:** 10.1101/2023.01.13.523917

**Authors:** Li He, Jiayu Wen, Qi Dai

**Affiliations:** Department of Molecular Bioscience, the Wenner-Gren Institute, Stockholm University; Division of Genome Sciences and Cancer, The John Curtin School of Medical Research, The Australian National University and Australian Research Council Centre of Excellence for the Mathematical Analysis of Cellular Systems.

## Abstract

BMP signalling acts as an instructive cue in various developmental processes such as tissue patterning, stem cell proliferation, and differentiation. However, it is not fully understood how this signalling pathway generates different cell-specific outputs. Here we have identified PRDM16 as a key co-factor for BMP signalling. PRDM16 contributes to a repressive role of BMP signalling on neural stem cell (NSC) proliferation. We demonstrate that PRDM16 regulates the genomic distribution of BMP pathway transcription factors, the SMAD4/pSMAD complex, preventing the activation of cell proliferation genes. When *Prdm16* is lost, the SMAD complex relocates to nearby genomic regions, leading to abnormal upregulation of BMP target genes. This function of PRDM16 is also required for the specification of choroid plexus (ChP) epithelial cells. Through a single-cell resolution fluorescent *in situ* approach, we have observed that genes co-repressed by SMAD and PRDM16, such as *Wnt7b* and several cell cycle regulators, become overexpressed in *Prdm16* mutant ChP. Our findings elucidate a mechanism through which SMAD4 and pSMAD1/5/8 repress gene expression. Moreover, our study suggests a regulatory circuit composed of BMP and Wnt signaling, along with PRDM16, in controlling stem cell behaviors.

## Introduction

Uncontrollable cell proliferation can lead to tumor growth, while premature differentiation can result in tissue degeneration. The balance between stem cell proliferation and differentiation is a crucial aspect in developmental and stem cell biology. BMP (Bone morphogenetic proteins) signaling is a key cell-signaling pathway in regulating stem cell proliferation and maintaining adult stem cell quiescence ^1–3^. Moreover, BMP signaling is essential in various cell specification processes ^4–6^.

The ability of a single pathway to play a diverse range of roles relies on context-specific transcriptional outputs. BMPs signal through two types of SMAD proteins: the receptor SMADs (R-SMADs) - Smad1, 5 and 8, and the co-SMAD protein SMAD4. Upon ligand binding, heterodimeric receptors like BMPRI and BMPRII phosphorylate R-SMADs, leading to the assembly and nuclear translocation of the SMAD complex with SMAD4. This complex then regulates gene expression by binding to enhancers of BMP target genes. Apart from transducing BMP signaling, SMAD4 is an essential effector in TGF-β/Activin signaling. In response to ligands such as TGF-β and Activin, SMAD4 associates with two other R-SMADs - phosphorylated SMAD 2 and 3, regulating downstream genes of the TGF-β/Activin signaling pathway. These SMADs recognize and directly bind to two main types of short DNA motifs in target enhancers via their N-terminal MH1 domain ^7^. However, since the binding is generally weak, SMAD complexes rely on various co-factors such as transcription factors, co-activators and co-repressors to activate or repress target gene expression ^8^.

In the mammalian brain, BMP and Wnt signaling pattern the brain midline where an essential structure, the ChP, emerges. Neural epithelial cells at the presumptive ChP site lose neural potential and exit cell cycle. Only a small number of cells at the boarder of the ChP primordium and cortical hem (CH) persist as slowly dividing ChP progenitors, leading to the expansion of the monolayered ChP epithelial cells ^9^. BMP signaling is essential for ChP epithelium specification. Conditional depletion of the BMP receptor *BMPr1a* diminishes ChP development ^10^, whereas ectopic BMP transforms neural progenitors into ChP cells ^11^. Wnt signaling, peaking at CH, is also necessary for ChP epithelial cell specification ^12^. Conditional depletion of ß-Catenin results in defective ChP, and overexpression of ß-Catenin expands CH at the expense of the ChP epithelium. These findings emphasize the importance of tightly controlling Wnt and BMP signaling levels for proper cell specification.

The PR domain-containing (PRDM) family protein PRDM16 determines cell fate specification in various cell types ^13–19^. Previous studies, including our own, demonstrated that *Prdm16* knockout (KO) mouse brains show severely reduced ChP structures ^20–22^. Interestingly, it was shown that PRDM16 can interact with TGF-ß pathway SMAD proteins *in vitro* and impact TGF-ß signaling output in craniofacial tissues ^21,23^. PRDM16 is also a downstream effector of BMP signaling during brown adipocyte specification ^24,25^. A recent study reported that PRDM16 and its ortholog PRDM3 (also known as Evi1) regulate Wnt signaling during craniofacial development in zebrafish ^26^. These findings suggest that PRDM16 may be more broadly involved in regulating BMP/TGF-ß and Wnt signaling. However, the underlying molecular mechanisms remain unclear.

Consistent with its essential roles in normal development, mutations and dysregulation of *Prdm16* are linked with several human diseases, including those identified in the patients with 1p36 chromosomal aberrations and cardiomyopathy ^27^. PRDM16 exhibits versatile functions at the molecular level, regulating chromatin accessibility and epigenetic states of its bound enhancers ^18,20^. Depending on associated cofactors, PRDM16 can either repress or activate gene expression. This dual role poses a challenge when considering PRDM16 as a therapeutic target, as undesired outcome may occur. Thus, a comprehensive understanding of the regulatory roles of this protein in each specific process is crucial for effective disease treatment strategies.

In this study we have investigated the mechanisms that regulate the transition between stem cell proliferation and differentiation during ChP development. We show that *Prdm16* mutant ChP cells fail to exit cell cycle, a similar phenotype to when BMP signaling is abolished ^10^. Using primary NSC culture, we dissected the molecular interaction of SMAD4/pSMAD1/5/8 proteins with PRDM16, and found that PRDM16 functions as a co-repressor that holds the SMAD complex at distal enhancers, repressing genes involved in cell proliferation. Finally, we validated that some of the co-regulated genes by BMP signaling and PRDM16 become derepressed in the *Prdm16* mutant ChP. These findings uncover an essential function of PRDM16 in stem cell regulation and BMP and Wnt signalling.

## Results

### PRDM16 promotes cell cycle exiting of neural epithelial cells at the ChP primordium

To understand how *Prdm16* regulates ChP epithelial specification, we first investigated the expression of this gene in the developing mouse brain. At embryonic day 10.5 (E10.5), BMP signaling specifies the presumptive ChP ^10^. *Prdm16* mRNAs and nuclear localization of the PRDM16 protein become detectable in the presumptive ChP at this stage (**Fig. 1A, supplementary Fig. 1A**). As previously shown, *Prdm16^cGT^*, a *Prdm16* knockout allele (*Prdm16 KO*), displayed severely reduced ChP structure at E13.5 (**Fig 1B)** ^20–22^. Expression of *Ttr*, a classic ChP marker gene, is reduced in both the lateral ventricle (tChPs) (**Fig. 1C**) and the fourth ventricle (hChP) (**Supplementary Fig. 1B**) in the *Prdm16 KO* mutant, suggesting that the function of PRDM16 is not area-restricted but generally required for ChP development.

**Figure 1.**
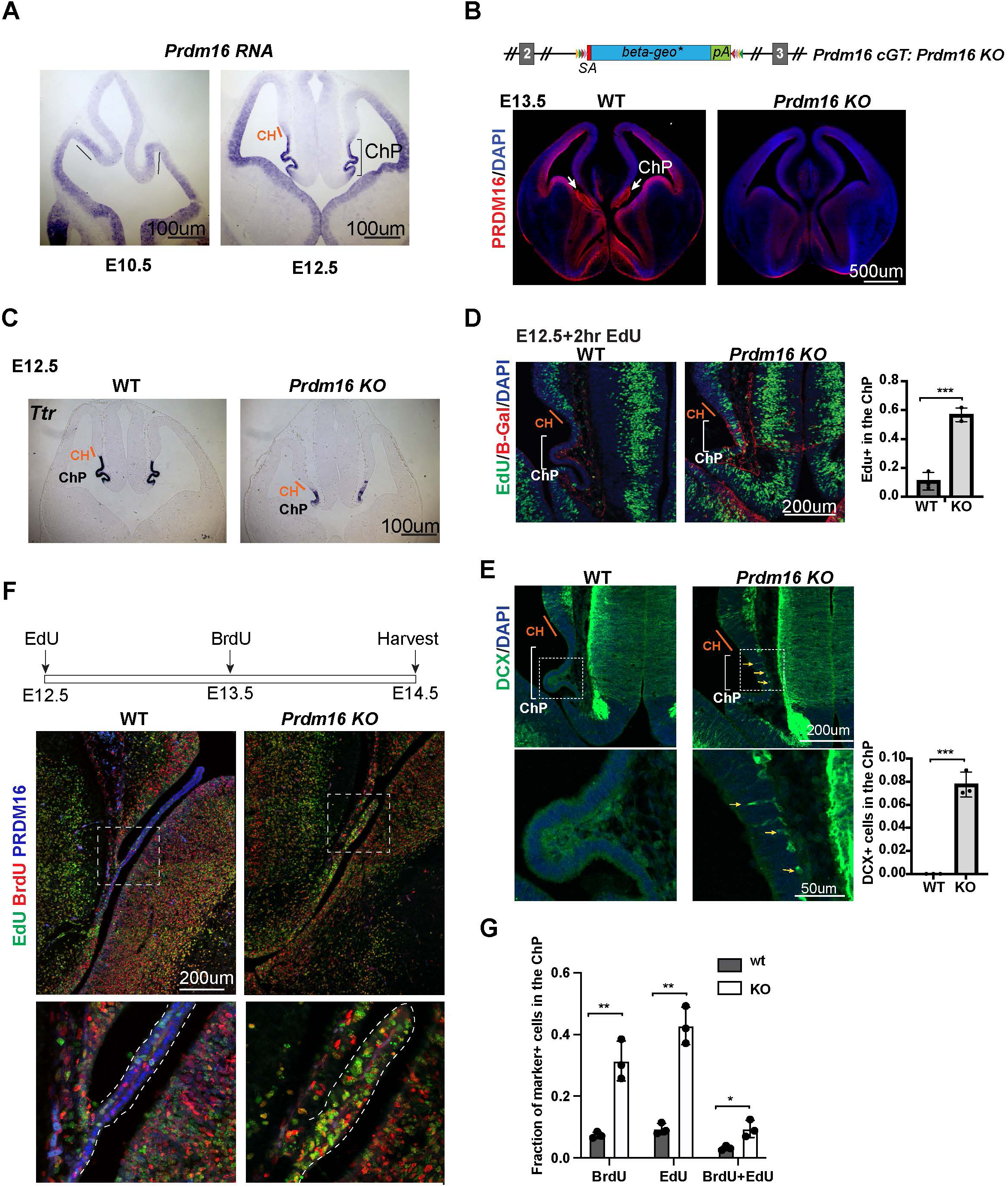
*Prdm16* mutant ChP cells fail to exit the cell cycle and retain neural progenitor identity. (**A**) RNA *in situ* hybridization using a *Prdm16* probe shows *Prdm16* expression in E10.5 and E12.5 brains. The line and bracket mark the developing CH and ChP. (**B**) Top: Schematic of the *Prdm16* GT allele, as reported in Strassman A. et al, 2017. Bottom: immunostaining of PRDM16 in E13.5 control and null mutant brains. Red: PRDM16; blue: DAPI. White arrows indicate the ChP in the control brain. (**C**) RNA in situ hybridization for *Ttr* on E12.5 control and *Prdm16* null mutant forebrains. Cortical hem (CH) regions are outlined based on morphology. (**D**) Two-hour EdU labeling in E12.5 control and *Prdm16* homozygous brains. Sections are co-stained with anti-β*-*Gal antibody to identify mutant ChP cells in the KO brain. Note: the β*-Geo* trap is absent in wild-type animals. Brackets mark ChP regions. Red: β-Gal; blue: DAPI; green: EdU. (**E**) Immunostaining for DCX on E12.5 control and *Prdm16* homozygous brain slices. Bottom panels show magnified views of the ChP (dashed rectangles). Yellow arrows indicate some of the DCX-positive cells. Right panels show quantification of EdU+ (**D**) and DCX+ cells (**E**). (**F**) Schematic of the double S-phase labeling experiment (top). Brain sections from E14.5 control and prdm16 KO embryos were immunostained with PRDM16 and BrdU antibodies and processed for EdU detection. Bottom panels show magnified views of the ChP region. (**G**) Quantification of EdU+ and BrdU+ cells from the double labeling experiment. Biological replicates: N=3. Error bars represent standard deviation (SD). Statistical significance is calculated using unpaired t-test. ***, p<0.001.

To assess the patterning of the CH and ChP regions, we analyzed the expression of *Wnt2b* and *BMP4* using conventional RNA *in situ* hybridization. *Wnt2b* expression, which marks the CH, appeared comparable between *Prdm16* KO and control brains at E11.5 and E12.5 (**Supplementary Fig. 1C**), indicating that CH development is not affected by the loss of *Prdm16*. BMP4, which labels the ChP and CH, also showed normal expression levels and spatial distribution in the mutant brain. These findings suggest that the BMP-dependent patterning of the ChP and CH domain remains intact in the absence of *Prdm16*.

However, the ChP epithelial layer in *Prdm16* mutants remained abnormally thick, resembling the adjacent neural epithelium (**Fig. 1C, Supplementary Fig. 1C**). This observation led us to investigate the underlying cellular changes responsible for the ChP defects in the mutant. To assess cell proliferation, we performed a 2-hour EdU labeling at E12.5. In control embryos, ChP cells were largely EdU negative and formed a monolayer, indicating that most had exited the cell cycle. In contrast, *Prdm16* mutant ChP cells marked by β*-Gal* remained highly proliferative (**Fig. 1D**). These results suggest several possibilities: ChP epithelial cells is not properly specified in the mutant, mutant ChP cells are specified but fail to exit the cell cycle to differentiate, or both processes are impaired.

To explore these possibilities, we first examined the cell type composition of *Prdm16* mutant ChP cells at E12.5. We stained brain sections from control and mutant embryos with the neuronal marker Doublecortin (Dcx). In controls, Dcx-positive cells were absent from the ChP, indicating that neural epithelial cells had successfully transitioned into ChP epithelial cells and lost their neural potential (**Fig. 1E**). In contrast, the *Prdm16* mutant ChP exhibited numerous Dcx-positive cells along the basal side of the epithelium, suggesting ongoing neurogenesis.

We next examined the expression of the NSC marker SOX2. In control brains, SOX2 is highly expressed in NSCs adjacent to the ChP epithelium but is significantly downregulated within the presumptive ChP region (**Supplementary Fig. 1D**). This downregulation was absent in the *Prdm16* mutant. Together, these results indicate that at E12.5, *Prdm16* depletion disrupts the normal transition of neural progenitors into ChP epithelial cells, maintaining cells in a proliferative, undifferentiated state.

We then asked whether PRDM16 not only regulates ChP epithelial specification but also directly restricts cell proliferation. To test this, we examined *Prdm16* mutant ChP cells at a later stage. We performed double S-phase labeling by injecting EdU at E12.5 and BrdU at E13.5, followed by sample collection at E14.5. By this stage, the wild-type ChP had developed into an elongated, monolayered epithelium in which most cells were negative for both EdU and BrdU, indicating cell cycle exit. In contrast, the *Prdm16* mutant ChP remained a disorganized cluster of cells, many of which were EdU-positive, BrdU-positive, or both (**Fig. 1F-G**). This finding suggests that although the mutant ChP can separate from the neuroepithelium at later stages, it fails to exit the cell cycle and differentiate, supporting a direct role of PRDM16 in restricting proliferation.

Given that BMP signaling is a key regulator of ChP formation and that the BMP pathway mutants such as *Foxg1-cre*::loxp*Bmpr1a* also exhibit ectopic cell proliferation in the developing ChP ^10^, we next investigated whether PRDM16 interacts with the BMP pathway to control cell proliferation.

### PRDM16 and BMP signaling collaborate to induce NSC quiescence *in vitro*

To investigate the molecular interplay between PRDM16 and BMP signaling in regulating cell proliferation, we turned to primary NSCs as an *in vitro* model. BMP signaling is known to maintain quiescence in adult NSC *in vivo* and can induce proliferative NSCs into quiescence *in vitro* ^1^. Furthermore, previous studies have shown that embryonic cortical NSCs are responsive to BMP4 treatment ^28^, making them a suitable system for probing downstream BMP signaling events. Based on this, we established a cell culture assay to evaluate the effects of BMP4 and PRDM16 on NSC proliferation and quiescence (**Fig. 2A**).

**Figure 2.**
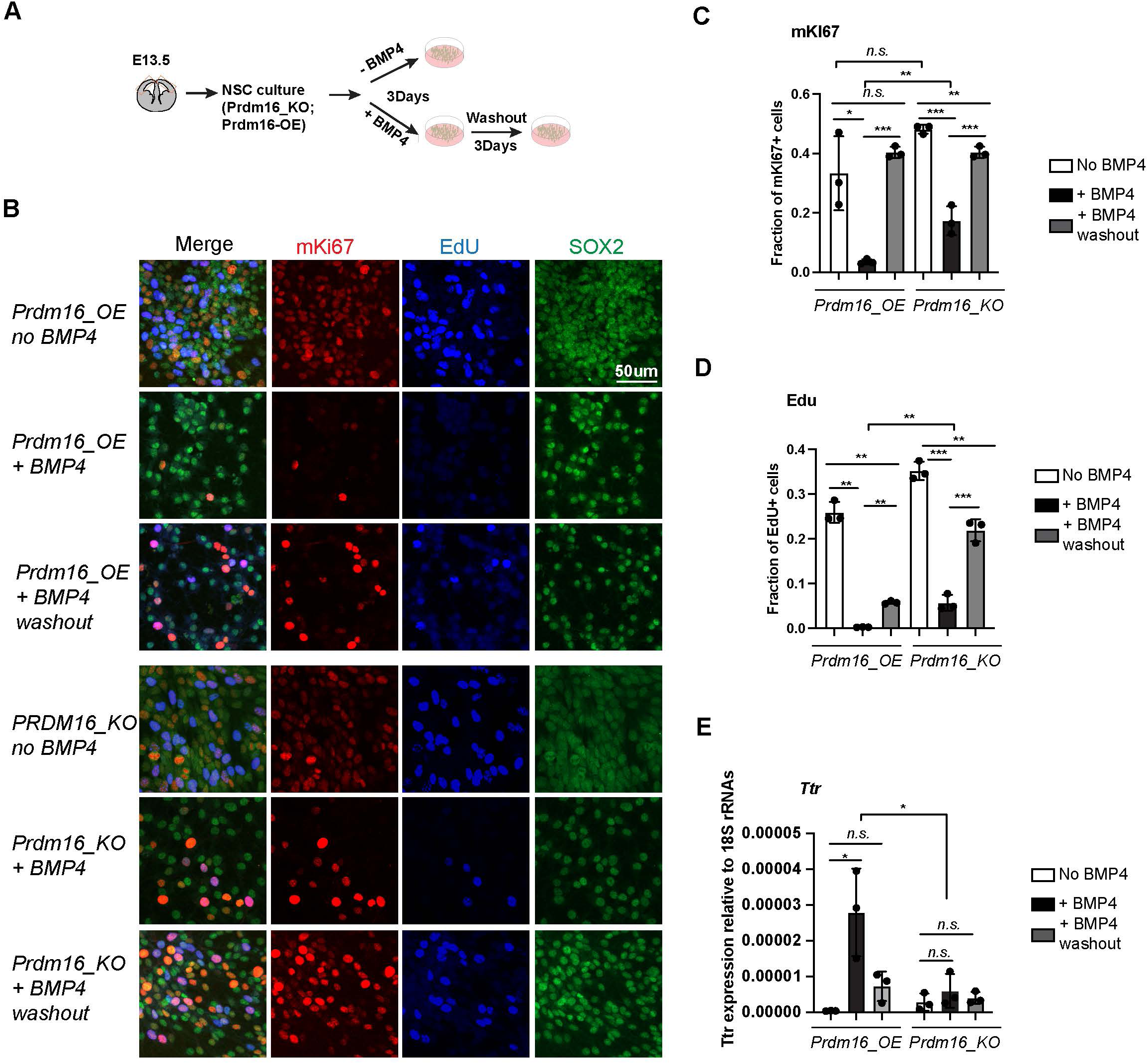
*Prdm16* is required for BMP4-induced NSC quiescence. (**A**) Schematic of the NSC culture assay. (**B**) Immunostaining of mKI67 in red and SOX2 in green and EdU labeling in blue on indicated NSC genotypes and treatment. (**C-D**) Quantification of the fraction of mKI67+ (C) and EdU+ (D) cells among the total cell number marked by SOX2. (**E**) RT-qPCR measurement of *Ttr* levels relative to 18S RNAs. Biological replicates: N=3. Error bars represent standard deviation (SD). Statistical significance is calculated using unpaired t-test. ***, p<0.001; **, p< 001; *, p<0.05; n.s., non-significant.

Unexpectedly, we found that PRDM16 protein levels were undetectable in cultured NSCs despite high levels of *Prdm16* mRNAs (**Supplementary Fig. 2A-B**). Given that the PRDM16 protein is normally restricted to the nucleus of NSCs in embryonic brain tissues (**Fig. 1B**, **Supplementary Fig. 1A** and ^20^), this observation suggests that *in vivo* mechanisms may regulate PRDM16 protein stability and nuclear localization. Post-translational modification of PRDM16 has been identified as a key regulatory mechanism in brown adipocytes ^29^, and we speculate a similar mechanism may operate in NSCs.

To bypass this limitation and examine PRDM16’s molecular function in regulating NSC proliferation and gene regulation, we generated a lenti-viral construct expressing *3xNSL_Flag*_*Prdm16* under a constitutive promoter. Infection of wild-type primary NSCs with this construct yielded a cell line with robust *Prdm16* mRNA expression and detectable nuclear PRDM16 protein (**Supplementary Fig. 2A-B**), which we referred to as *Prdm16_overexpressing (Prdm16_OE)*. For comparison, we attempted to establish three additional lines: wild-type NSCs infected with the empty vector (*wt_CDH*), *Prdm16_KO* NSCs infected with the empty vector (KO_CDH), and *Prdm16_KO* NSCs infected with *3xNSL_Flag*_*Prdm16.* However, we were unable to generate the last line despite repeated attempts. This failure was likely due to low viral production (the *Prdm16* coding sequence exceeds 3kb) and the increased sensitivity of KO NSCs to viral infection. Nevertheless, we proceeded with comparative analyses of *Prdm16_OE* cells against *wt_CDH*, *KO_CDH* and uninfected *KO* NSCs.

To assess cell proliferation rates, we labeled NSCs with EdU and mKi67. BMP4 treatment of *Prdm16_OE* NSCs led to a marked reduction in the number of mKi67- and EdU-positive populations (**Fig. 2B-D**). Following BMP4 washout, *Prdm16_OE* NSCs restored EdU and mKi67 labeling, indicating re-entry into the cell cycle. This reversible reduction confirms that BMP4 induces a quiescent, non-proliferative state in NSCs. In contrast, a larger proportion of *Prdm16*_KO cells failed to exit the cell cycle in response to BMP4, as shown by a less pronounced decrease in mKi67- and EdU-positive populations (**Fig. 2B-D**). Notably, *Prdm16_KO* cells exhibited similar properties regardless of whether they were infected with the control viral vector (**Supplementary Fig. 2D-F)**, and we therefore used both lines interchangeably in our analyses.

Previous studies have shown that BMP4 can program ChP cell fate and activate ChP-specific genes in neural progenitors cultured *in vitro* ^11^. To test whether PRDM16 cooperates with BMP signaling to induce ChP gene expression, we measured *Ttr* mRNA levels using reverse-transcription followed by quantitative PCR (RT-qPCR). In *Prdm16_OE* NSCs, *Ttr* expression was robustly induced following BMP4 treatment. In contrast, *Prdm16*_KO NSCs and NSCs lacking the *Prdm16*_OE construct failed to upregulate *Ttr* in response to BMP4 (**Fig. 2E, Supplementary Fig. 2G**).

Together, these results indicate that both BMP signaling and PRDM16 are required not only to restrict NSC proliferation but also to induce ChP gene expression. We next investigated the molecular mechanisms underlying this cooperation.

### BMP signaling and PRDM16 cooperatively repress proliferation genes

To understand how BMP signaling and PRDM16 suppress cell proliferation, we aimed to determine the transcriptional targets of SMAD4 and pSMAD1/5/8 in cells with and without BMP4. We first applied Cleavage Under Targets and Tagmentation (CUT&TAG) experiments using a PRDM16 antibody and the available SMAD antibodies to profile their genomic binding sites. The PRDM16 antibody worked efficiently, but none of the SMAD antibodies produced libraries with sufficient sequencing reads. Subsequently we employed chromatin immunoprecipitation followed by deep-sequencing (ChIP-seq) for the SMAD proteins. Given that SMAD4 forms a complex with SMAD2/3 only in response to TGF-β/activin-type ligands, we included an antibody targeting SMAD3 as a control for non-BMP4-induced SMAD4 binding. Thus, our experiment settings enabled us to profile genomic binding sites for PRDM16, SMAD4 and two types of R_SMADs under conditions with endogenous BMPs and TGF-β/Activin, as well as ectopic BMP4.

In *Prdm16_OE* cells without BMP4, we identified several hundred to a few thousand ChIP-seq peaks for all three classes of SMAD proteins after normalizing to the input reads (FDR < 10%), indicating endogenous levels of BMP and TGF-β signaling in these cells (**Fig. 3A, Supplementary Fig. 3A-C**). Following BMP4 addition, pSMAD1/5/8 exhibited approximately a 750-fold increase in peak number, while the number of SMAD3 peaks increased by less than threefold (**Fig. 3A**, **Supplementary Fig. 3A**). Furthermore, in cells treated with BMP4, the pSMAD1/5/8 peaks largely overlap with the SMAD4 peaks, but much less with the SMAD3 peaks (**Supplementary Fig. 3B**). For example, pSMAD1/5/8 binds to two enhancers in the intronic region of *Prdm16,* with the binding intensity dramatically increased in response to BMP4 (**Supplementary Fig. 3D**). This suggests that *Prdm16* itself is a transcriptional target of BMP signaling. Indeed, *Prdm16* mRNA levels were elevated by BMP4 in wild-type but not *Prdm16_KO* cells (**Supplementary Fig. 2B**). By contrast, SMAD3 showed little ChIP-seq signal in the *Prdm16* gene locus even in the presence of BMP4. These results confirm that the response of pSMAD1/5/8 is specific to BMP4, while SMAD3 is generally unresponsive to BMP4. The gained Smad3 sites likely result from an indirect effect of BMP signaling.

**Figure 3.**
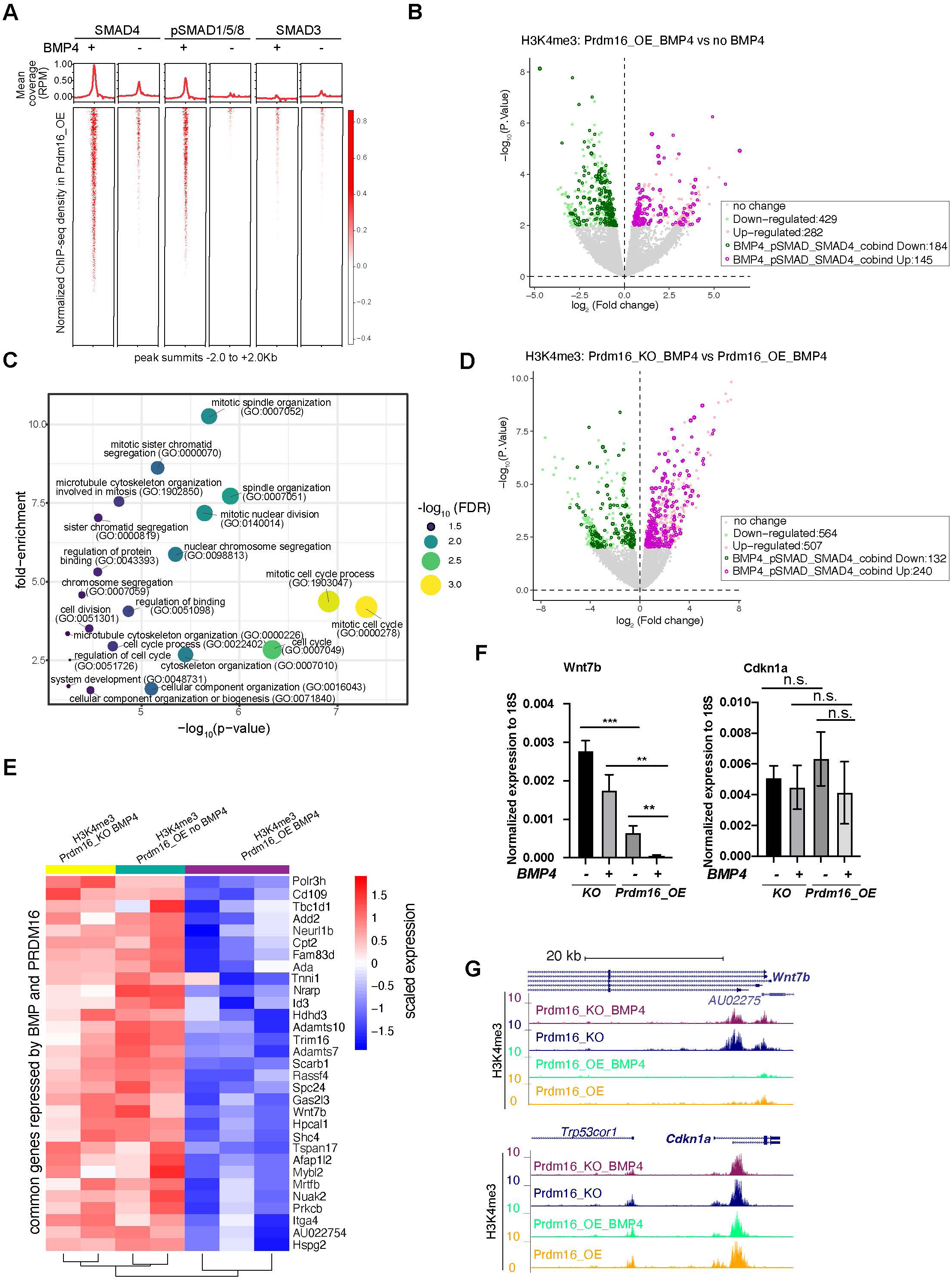
BMP signaling represses cell proliferation genes to induce cell quiescence. **(A)** Heatmaps of ChIP-seq coverage (normalized by library depths and input) centered at peak summits for SMAD4, pSMAD1/5/8 and SMAD3 in *Prdm16_OE* cells with or without BMP4. **(B)** Volcano plot displaying differential H3K4me3 signal at TSS in *Prdm16_OE* cells with BMP4 versus untreated controls. Genes with significant changes (*P* < 0.01) are colored. (**C**) Bubble plot of GO terms significantly enriched among SMAD-repressed genes (highlighted in dark green in panel B). (**D**) Volcano plot of differential H3K4me3 signal at TSS comparing *Prdm16_KO* and *Prdm16_OE* cells, both treated with BMP4. Significantly changed genes (*P* < 0.01) are colored. **(E)** Heatmap of normalized H3K4me3 read counts at the TSS of 31 genes co-repressed by PRDM16 and SMADs (overlapping genes from dark green in B and dark pink in D). (**F**) RT-qPCR quantification of *Wnt7b* and *Cdkn1a* mRNA levels across indicated genotypes and treatment conditions. Data represent three biological replicates, each with two technical replicates. Statistical significance was calculated using unpaired t-test (***, p<0.001; **, p< 0.01; *, p<0.05; n.s., non-significant). (**G**) Genome browser tracks of H3K4me3 CUT&TAG profiles at the *Wnt7b* and *Cdkn1a* loci.

To evaluate the regulatory activity of SMAD4/pSMAD1/5/8 in NSCs, we determined the state of transcription activation using H3K4me3 CUT&TAG signals at annotated gene transcription start sites (TSS) in BMP4-treated and untreated *Prdm16_OE* cells. As cell identity genes are known to be marked by extended breadth of H3K4me3 ^30^, we reasoned that compared to RNA-seq, which measures all gene products, changes in the H3K4me3 signal at the promoter might bias toward cell identity genes between proliferation and quiescence states. Thus, we determined all BMP4-repressed and activated genes by measuring changes in the H3K4me3 CUT&TAG read coverage at annotated TSS in BMP4-treated *Prdm16_OE* cells compared to untreated, identifying 282 up-regulated and 429 down-regulated genes (**Fig. 3B**) (P<0.01). To pinpoint genes directly regulated by SMAD4 and pSMAD1/5/8, we intersected the dysregulated genes with those whose regulatory regions contain overlapping SMAD4 and pSMAD1/5/8 ChIP-seq peaks. This analysis led to the identification of 145 up-regulated and 184 down-regulated SMAD4 and pSMAD1/5/8 target genes.

To elucidate the function of BMP-regulated targets, we performed gene ontology (GO) analyses for the genes that changed expression in response to BMP4 in *Prdm16_OE* cells and were also bound by SMAD4 and pSMAD1/5/8. Remarkably, the downregulated genes (BMP4-repressed genes), but not the up-regulated genes (BMP4-activated genes), showed significantly enriched functional categories, with nearly all GO terms related to cell proliferation/cell cycle (**Fig. 3C)**. A recent study reported that BMP2-induced genes are enriched for neuronal and astrocyte differentiation ^31^, while our analysis did not identify these categories. One possibility for the discrepancy is that we overexpress *Prdm16* in cultured NSCs, which may reinforce BMP signaling activities in cell proliferation. Other possibilities could be the use of different BMP ligands (BMP4 versus BMP2), differences in the origin of NSC culture (ours was from E13 instead of E11 and E14), or different profiling methods (H3K4me3 versus RNA-seq).

To identify PRDM16-repressed and activated genes under active BMP signaling conditions, we compared changes of H3K4me3 coverage at TSSs between BMP4-treated *Prdm16_KO* cells and BMP4-treated *Prdm16_OE* cells. We found approximately twice as many up-regulated genes (240) as down-regulated genes (132) (**Fig. 3D**), suggesting that the cooperative activity of PRDM16 and BMP signaling mainly represses gene expression. We further overlapped the 184 genes repressed by BMP4 and the 240 genes repressed by PRDM16, identifying 31 common ones. These 31 genes displayed low H3K4me3 coverage in *Prdm16_OE* cells with BMP4 but higher H3K4me3 signal in both *Prdm16_KO* cells treated with BMP4 and untreated *Prdm16_OE* cells (**Fig. 3E).**

Next, we attempted to validate whether changes in TSS H3K4me3 intensity correspond to changes in mRNA levels, by conducting RT-qPCR for selected genes whose function is associated with cell proliferation from the gene ontology analysis: *Wnt7b*, *Mybl2*, *Id3*, *Spc24* and the three *Spc24*-related genes (*Spc25*, *Ndc80* and *Nuf2*) (**Fig. 3F-G, Supplementary Fig. 3E-K**). SPC24 forms the NDC80 kinetochore complex along with three other proteins: SPC25, NDC80 and NUF2 ^32^ (Illustrated in **Supplementary Fig. 3H**), and their function is essential for chromosome segregation and spindle checkpoint activity. Notably, *Spc25*, *Ndc80* and *Nuf2* appeared repressed by PRDM16 and BMP4 based on changes in H3K4me3 at the TSS (**Supplementary Fig. 3E**), despite not being identified as the top 31 candidates. mRNA levels for most of the tested genes followed a similar pattern to the H3K4me3 intensity: BMP4-treated *Prdm16_OE* cells showed the lowest expression, while either the loss of *Prdm16* or absence of BMP4 led to upregulation. *Spc24* gene products showed minimal amplification in qPCR, likely due to poor primer sequence quality or unstable mRNAs. Another exception is *Id3* whose expression increased upon BMP4 treatment or *Prdm16* depletion, indicating PRDM16 repressing *Id3* but BMP4 inducing *Id3*. However, the activation of *Id3* by BMP4 is significantly stronger in *Prdm16_KO* cells than that in *Prdm16*_*OE* cells, suggesting that BMP signaling normally activates *Id3* but PRDM16 suppresses such activation. In contrast, *Cdkn1a,* a target gene of TGF-β pathway encoding the cell cycle inhibitor P21, exhibited consistent H3K4me3 coverage and mRNA levels across BMP4 treated and non-treated cells, as well as between *Prdm16_OE* and *KO* cells (**Fig. 3F-G**), confirming that neither PRDM16 nor BMP signaling influences *Cdkn1a* expression in NSCs.

### PRDM16 assists genomic binding of SMAD4 and pSMAD1/5/8

To understand how PRDM16 interacts with BMP signaling, we integrated PRDM16 CUT&TAG data with SMAD ChIP-seq data, focusing on assessing PRDM16’s influence on the genomic binding of SMAD proteins.

The SEACR peak caller software ^33^ identified 3337 and 7639 PRDM16 CUT&TAG common peaks from four replicates of BMP4-treated and untreated *Prdm16_OE* samples (FDR < 10%, see Methods), respectively (**Fig. 4A**). Upon BMP4 treatment, there were 5936 lost and 1634 gained PRDM16 CUT&TAG peaks, suggesting that PRDM16 regulates a distinct subset of genes in proliferating versus quiescent NSCs. Samples from *Prdm16*_*KO* cells lacked these sites, confirming the specificity of the PRDM16 CUT&TAG signal (**Fig. 4B**).

**Figure 4.**
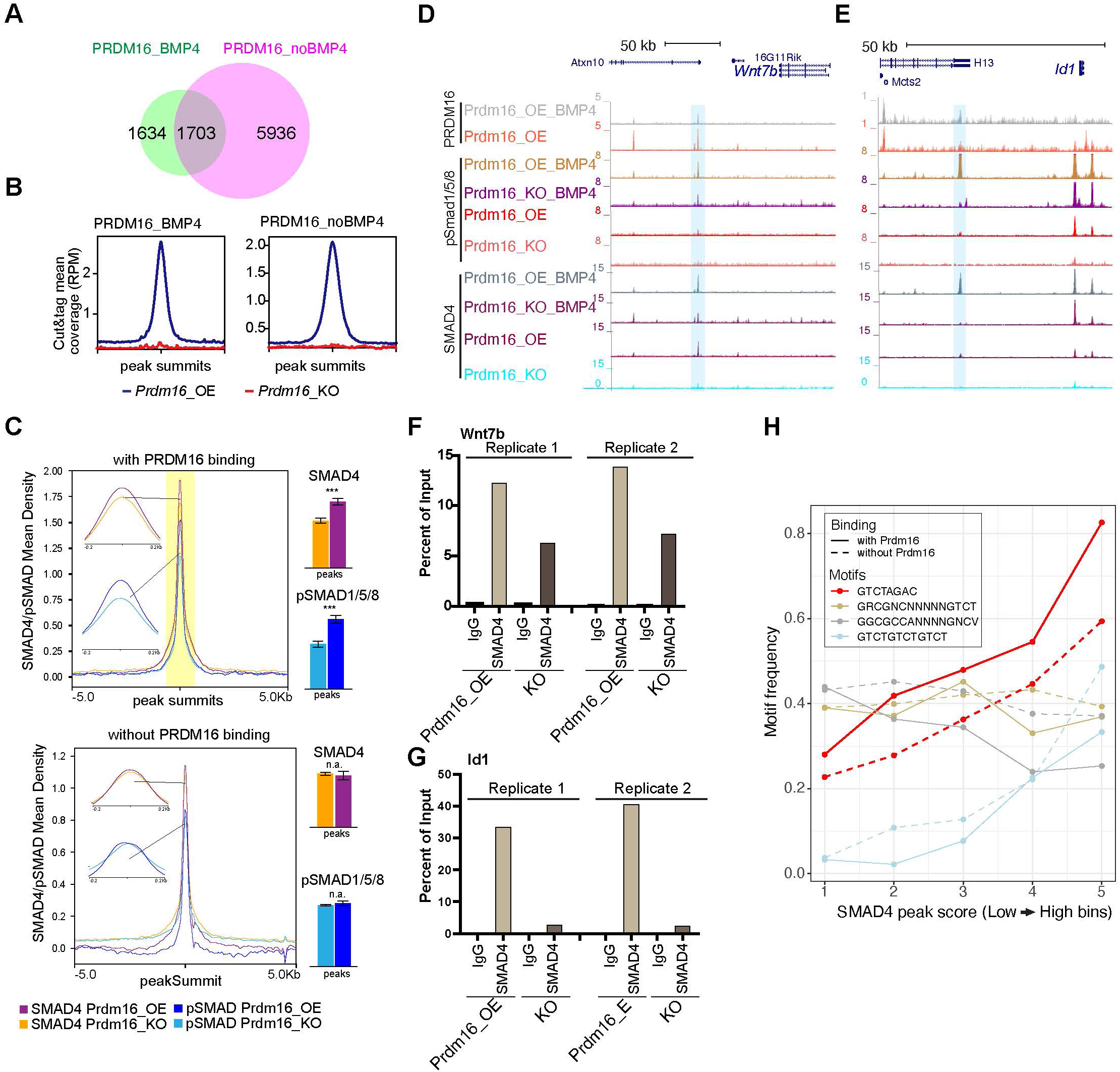
PRDM16 anchors SMAD proteins at specific genomic regions to mediate gene repression. (**A)** Venn diagram showing overlap of PRDM16 CUT&TAG peak in BMP4-treated and untreated *Prdm16_OE* cells. **(B)** Average PRDM16 CUT&Tag signal (normalized to library size) at peak summits in BMP4-treated, untreated Prdm16_OE, and Prdm16_KO cells (4 replicates combined). **(C)** Metaplots of SMAD4 and pSMAD1/5/8 ChIP-seq coverage centered on peak summits with (top) and without (bottom) PRDM16 co-binding in BMP4 treated *Prdm16_OE* and *Prdm16_KO* cells. Bar plots (right) show average coverage; significant reduction in the KO cells is seen only at PRDM16 co-bound sites (*** P<0.001; ** P<0.01; * P<0.05; n.a. P>0.05). **(D-E)** Genome browser view of *Wnt7b* and *Id1* loci showing PRDM16 CUT&TAG and SMAD ChIP-seq tracks. Co-bound regions are highlighted. **(F-G)** ChIP-qPCR validation of SMAD4 binding at highlighted regions in D-E from *Prdm16_OE* and *Prdm16_KO* cells (2 replicates). (**H**) Frequency of four SMAD motif types across SMAD ChP-seq peaks binned by binding strength. The palindromic motif GTCTAGAC is most enriched at PRDM16 co-bound sites (solid red) and notably reduced in non-co-bound peaks (dashed red).

From our overlapping analyses (see Methods), we found that over 50% of SMAD4 and pSMAD1/5/8 binding peaks were consistent in *Prdm16_OE* and *Prdm16_KO* cells (**Supplementary Fig. 4A-B**), indicating that deletion of *Prdm16* does not affect general genomic binding ability of these proteins. Using Homer’s mergePeaks function with PRDM16 CUT&TAG and SMAD ChIP-seq data, we identified co-bound sites by PRDM16 and SMAD proteins in BMP4-treated *Prdm16_OE* cells. PRDM16 CUT&TAG peaks mainly overlap with SMAD4 and pSMAD1/5/8 peaks, but not much with SMAD3 peaks (**Supplementary Fig. 4A-C**). This result suggests that PRDM16 mainly collaborates with the SMAD4/pSMAD1/5/8 complex but not the SMAD3/SMAD4 complex in cells with high levels of BMP4.

Further examination of SMAD4 and pSMAD1/5/8 binding revealed significantly lower enrichment of SMAD4 and pSMAD1/5/8 at PRDM16 co-bound sites in *Prdm16_KO* cells compared to *Prdm16_OE* cells (**Fig. 4C**) (P=2.5E-6, and P=4.7E-3, respectively, two-tailed *t-*test). As a control, the SMAD4 and pSMAD1/5/8 sites without PRDM16 co-binding did not show such change (P>0.05). This result suggests that SMAD binding to the PRDM16 co-binding sites depends on PRDM16.

To validate the co-binding of SMAD4 and PRDM16, we selected the candidate loci, *Wnt7b* and *Id1* (**Fig. 4D-E**) and applied sequential ChIP-qPCR. This experiment confirmed simultaneous binding of SMAD4 and PRDM16 to the same DNA molecules at these loci (**Supplementary Fig. 4D**), as SMAD4 was more enriched in the chromatin pulled by the PRDM16 antibody compared to the IgG control. Additionally, ChIP-qPCR confirmed reduced SMAD binding at the SMAD/PRDM16 co-bound site in *Prdm16_KO* cells compared to *Prdm16_OE* cells (**Fig. 4F-G**). Thus, PRDM16 enhances genomic binding of the SMAD proteins to specific genome regions.

### PRDM16 facilitates SMAD4 binding to regions enriched for SMAD palindromic motifs

We then wondered whether there is sequence feature that distinguishes the regions co-bound by SMADs and PRDM16 from those only bound by SMADs. To this end, we checked SMAD-bound regions with PRDM16 binding and those without for two types of SMAD4 binding motifs ^8^. Together with pSMAD1/5/8 or pSMAD3, SMAD4 may bind to a palindromic motif, GTCTAGAC or direct repeats of GTCT like GTCTGTCTGTCT ^34–36^; together with pSMAD1/5/8, SMAD4 associates with GC-rich SBEs (SMAD-binding elements) including GGCGCC-AN4-GNCV and GRCGNCNNNNNGTCT ^37–39^. We calculated occurrence frequency of each of these motifs in binned SMAD4 ChIP-seq peaks from *Prdm16_OE* cells treated with BMP4 (bin 1 to 5 with low to high peak scores, **Fig. 4H**). Interestingly, the palindromic motif is the most enriched one. Its occurrence frequency increases with peak scores, but becomes lower in the SMAD4 peaks absent for PRDM16 co-binding. The frequency of the GTCT-triplet motif also increases with higher SMAD4 peak scores, while there is no reduction in regions without PRDM16 binding. The two GC-rich motifs show a distinct trend: they are not as highly enriched, their frequencies do not increase with higher SMAD4 peak scores, and the occurrence frequency in PRDM16 co-bound regions is either comparable to or even lower than the regions without PRDM16 binding. Together, this result suggests that PRDM16 may separate the SMAD4/pSMAD1/5/8 proteins into two types of genomic regions, one enriched for the palindromic motif where PRDM16 is present and the other with the GC-rich motifs.

### SMAD4 and pSMAD1/5/8 switch genomic locations in the absence of PRDM16

Our careful inspection on the *Wnt7b* and *Id1* loci surprisingly revealed that there are multiple ectopic SMAD4 and pSMAD1/5/8 peaks in the *Prdm16* mutant sample (arrow-indicated peaks in **Fig. 5A-B**). We then globally assessed the extent of ectopic SMAD binding surrounding the SMAD/PRDM16 co-bound sites by quantifying differential SMAD binding intensity between *Prdm16_KO* and *Prdm16_OE* cells (**Fig. 5C**, FDR < 10%). In agreement with the metaplots (**Fig. 4C**), all of the SMAD/PRDM16 co-bound sites (blue dots) showed reduced ChIP-seq read coverage in *Prdm16_KO* cells. By contrast, the flanking regions within the 200kb range of a co-bound site (red dots) displayed a trend of increase of SMAD binding in *Prdm16_KO* cells. This result agrees with the aforementioned motif analysis that PRDM16 helps to localize the SMAD complex at specific genomic regions, and it also suggests that without *Prdm16*, the SMAD factors are partly redirected to neighboring genomic regions.

**Figure 5.**
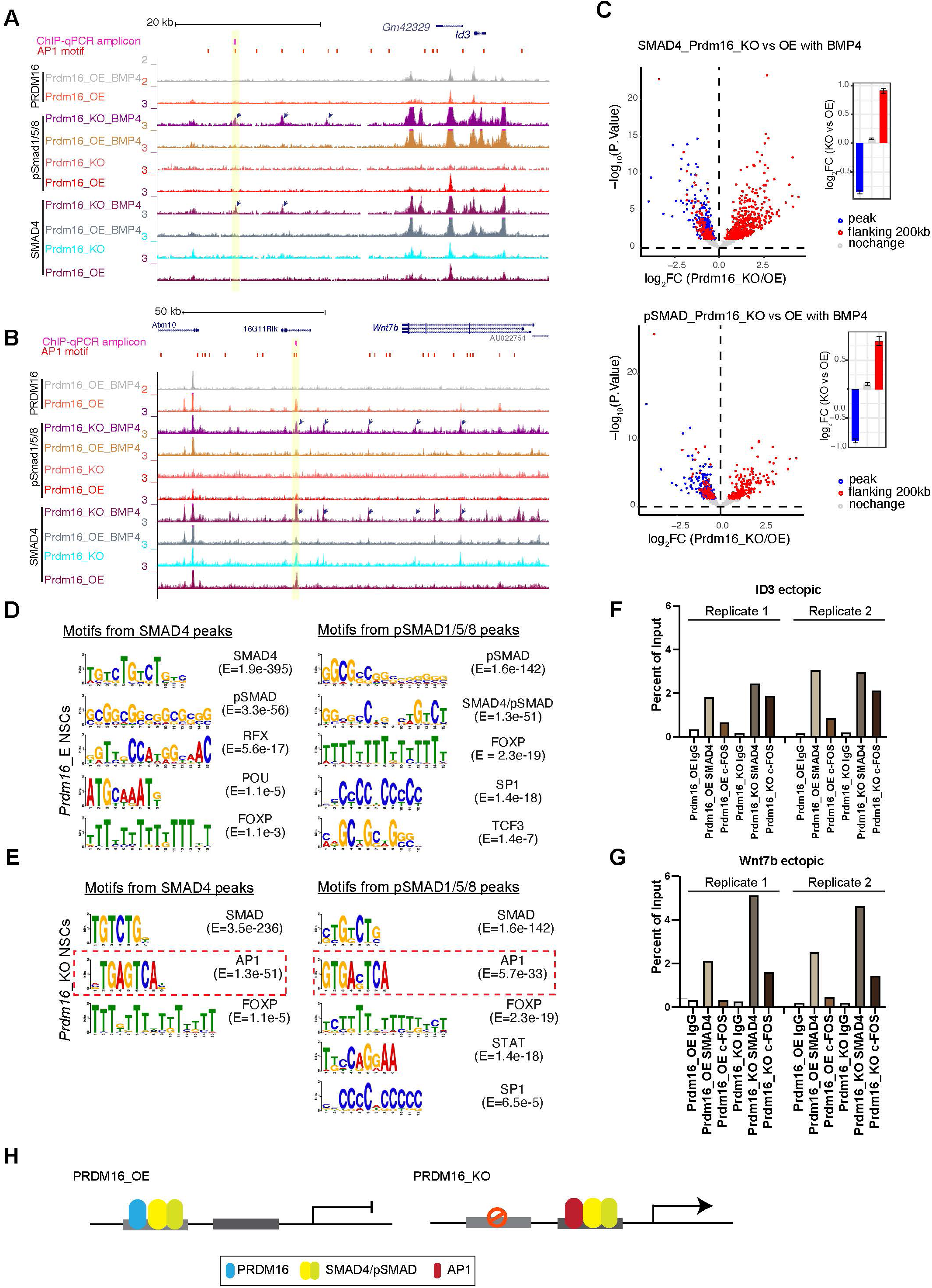
The SMAD complex shifts genome binding in the absence of Prdm16. (**A-B**) Genome browser views of the *Id3* (top) and *Wnt7b* (bottom) loci showing ectopic SMAD peaks (arrows) in *Prdm16 KO* cells. **(C)** Volcano plots showing differential ChIP-seq signal for SMAD4 and pSMAD1/5/8 in BMP4-treated *Prdm16_KO* cells versus BMP4-treated *Prdm16_OE* cells. Blue dots represent sites co-bound by PRDM16; red dots indicate sites not bound by PRDM16 but located within 200kb of a PRDM16-cobound region. Mean log2 fold-change values are plotted to the right. (**D-E**) *De novo* motif discovery from SMAD4 and pSMAD1/5/8 ChIP-seq peaks in *Prdm16_OE* cells (**D**) and *Prdm16_KO* cells (**E**). (**F-G**) ChIP-qPCR validation of increased SMAD4 and c-FOS occupancy at the indicated ectopic binding sites in *Prdm16_KO* versus *Prdm16_OE* cells. Locations of PCR amplicons are shown in A and B. **(H)** Model illustrating enhancer switching: in the absence of PRDM16, SMAD complexes alter their co-factors and function as transcriptional activators.

### AP1 is the potential co-factor that interacts with relocated SMAD proteins

We intended to find out how SMAD4 and pSMAD1/5/8 relocate to ectopic sites in the absence of *Prdm16*. By running *de novo* motif discovery (see Methods), we identified a number of significantly enriched DNA motifs from pSMAD1/5/8 and SMAD4 ChIP-seq peaks in *Prdm16_OE* and *Prdm16_KO* cells (**Fig. 5D-E**). Interestingly, in addition to the known SMAD motifs, the only other motif identified in both pSMAD1/5/8 and SMAD4 peaks from *Prdm16_KO* cells but not *Prdm16_OE* cells is the AP1 motif. Furthermore, the AP1 motif is more frequently found in the pSMAD1/5/8 and SMAD4 regions lacking PRDM16 binding than those with PRDM16 binding (**Supplementary Fig. 4E**). The interaction between the AP-1 complex with the SMAD proteins was reported previously ^40^. Our result implies that AP-1 is a potential cofactor for the SMAD proteins in the absence of *Prdm16*. To validate this finding, we performed ChIP-qPCR with an antibody against c-FOS, one of the subunits of AP1, on *Prdm16_OE* and *KO* cells. Supporting the global analyses, c-FOS is enriched in the Smad4- bound regions at the *Wnt7b* and *Id3* loci in the *Prdm16_KO* condition (**Fig. 5F-G**). There is little c-FOS binding to these regions in cells expressing *Prdm16*, suggesting that there is cooperativity of SMAD and AP-1 which facilitates each other’s binding when PRDM16 is absent. Alternatively, PRDM16 may suppress genomic accessibility of these regions for SMAD and AP-1 proteins, a function we reported for PRDM16 in cortical NSCs ^20^.

Together, our genomic data and validation suggest an enhancer switch model (**Fig. 5H)**: PRDM16 assists SMAD protein binding to repressive cis-regulatory elements that are enriched for GTCTAGAC palindromic motifs; loss of PRDM16 results in genomic relocation of SMAD proteins, presumably via the association with coactivators such as AP1; the consequence of SMAD relocation is de-repression of cell proliferation regulators.

### SMADs and PRDM16 co-repressed genes in NSCs are suppressed in the developing ChP

Next, we investigated whether genes co-regulated by PRDM16 and BMP signaling in NSCs (**Fig. 3C**) are also involved in ChP development, where both factors limit NSC proliferation ( ^10^ and **Fig. 1**). Using published E12.5 mouse brain single cell RNA-seq (scRNA-seq) data ^41^, we analyzed expression of the 31 co-repressed genes across ChP epithelial (*Ttr*+), CH (BMP4+/*Ttr*-) and a few forebrain-specific radial glia (RG) (Sox2+/Hes5+) clusters (**Fig. 6A, Supplementary Fig. 5A**). Except for one gene *Mrtfb* that was not found in the scRNA-seq data, thirteen showed no or weak expression (**Supplementary Fig. 5A**), and seventeen had moderate to high expression in at least one cluster (**Fig. 6A**). Notably, these genes were generally expressed at lower levels in ChP clusters than in CH or RG clusters, suggesting repression in the ChP.

**Figure 6.**
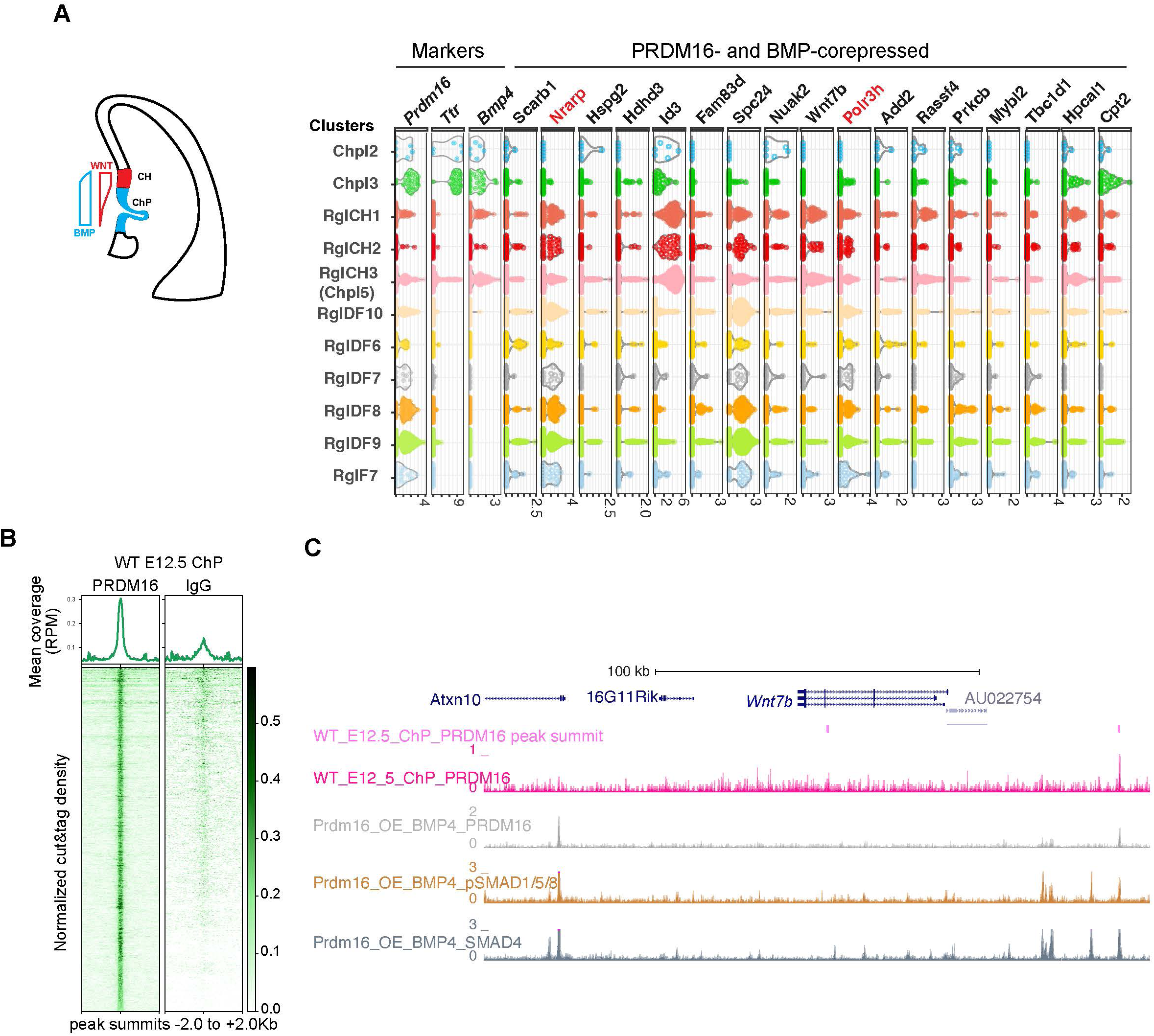
PRDM16 and SMADs co-regulated genes exhibit differential expressed in ChP and neural cells. (**A**) Schematic illustrating the CH and ChP regions, which exhibit opposing Wnt and BMP signaling gradients. Shown alongside is a violin plot of scRNA-seq expression at E12.5 for the indicated genes across cell type clusters ^37^. Cell clusters were defined based on marker gene expression from E9.5-E18.5 brain tissues, including choroid plexus clusters 1-5 (Chpl), with Chpl5 newly identified in this study based on *Ttr* expression as well as radial glia clusters: RglCH 1-3, RglDF 6-10, and RglF7. CH: cortical hem; DF: dorsal forebrain; F: forebrain. Gene expression per cell is represented as log2-normalized counts. **(B)** Metaplots and heatmaps showing CUT&TAG signal for PRDM16 and IgG, centered on peak summits and normalized for library size, from E12.5 dorsal midline tissues. **(C)** Genome browser view of the *Wnt7b* locus, displaying CUT&TAG tracks for PRDM16 and ChIP-seq tracks for SMAD proteins.

To test for direct PRDM16 regulation, we performed CUT&TAG assays using the PRDM16 antibody on dissected dorsal midline tissues that mainly contain CH and the ChP (**Fig 6B** and **Supplementary Fig. 5B**) and identified 3238 common peaks across three replicates. Manual inspection of the 31 loci revealed PRDM16 binding near the TSS of 25 genes using the browser Embryonic Mouse Brain Epigenome Atlas ^42^ (Text in black in **Fig. 6A** and **Supplementary Fig. 5A**). Using an unbiased peak-to-promoter mapping, 24/31 co-repressed and 84/153 BMP-only-repressed genes had PRDM16 binding in E12.5 ChP. The enrichment in the co-repressed group (**Supplementary Fig. 5C,** Fisher’s Exact Test, p = 0.015) indicates a stronger regulatory role of PRDM16 on these genes in the developing ChP.

As an example, *Wnt7b* has two PRDM16 peaks (**Fig 6C**), one unique to the dorsal midline tissue. Given Wnt signaling’s role in ChP specification and NSC proliferation^12,43^. *Wnt7b* likely represents a direct PRDM16 target in the developing ChP.

### PRDM16 represses *Wnt7b* and Wnt activity in the developing ChP

We sought to determine which of the identified genes from NSC culture are regulated by PRDM16 in the developing ChP. In addition to *Wnt7b*, several other Wnt ligands are expressed in the CH and ChP region ^44^. For example, *Wnt3a* is expressed in the dorsal-midline region but not in the forebrain, which explains why it was not identified from the forebrain-derived NSC datasets. Although there is no PRDM16 CUT&TAG peak within the *Wnt3a* locus in cultured NSCs, a PRDM16 CUT&TAG peak is present in the intronic region of *Wnt3a* in the dorsal midline tissue (**Supplementary Fig. 5D**). We thus examined the scRNA-seq data ^41^ and confirmed that multiple components in the Wnt and BMP pathways are present in the CH and ChP clusters (**Supplementary Fig. 5E**). To systematically measure expression changes of PRDM16 target genes in the *Prdm16* mutant brain, we applied a multiplexed fluorescent *in situ* approach, Single-Cell Resolution in Situ Hybridization On Tissues (SCRINSHOT) ^45^. In this method each mRNA molecule is hybridized with three gene-specific padlock probes and visualized by florescent dye-conjugated detection probes. A dot of fluorescent signal corresponds to one mRNA molecule, allowing quantitative measurement of gene expression in single cell resolution.

We designed probes for ten Wnt and five BMP pathway components, other genes that are co-repressed by BMP and PRDM16 in NSC culture, as well as cell-type specific markers (the ChP marker gene *Ttr* and *Foxj1*, the neural progenitor markers *Sox2*, *Hes5* and *Zbtb20,* and the neuronal marker Ngn2), and conducted SCRINSHOT in three pairs of control and *Prdm16* mutant brains.

In the control brain, the *Ttr* and Foxj1 probes detect the ChP epithelium but not the adjacent tissues at E11.5 and E12.5, confirming the signal specificity of SCRINSHOT. Several Wnt (*Wnt2b*, *Wnt7b*, *Wnt5a*, *Wnt3a*, *Fzd10* and *Axin2*) and BMP components (*BMP4* and *BMP7*) also display tissue specificity: the Wnt genes are more enriched at CH while the BMP genes are more at the ChP. The representative images from E11.5 and E12.5 samples are presented in **Fig. 7A-B**. Except *Wnt2b*, the other tested *Wnt* genes exhibit low expression in the ChP cells, consistent with the notion that Wnt signaling is present and required at the ChP ^12^.

**Figure 7.**
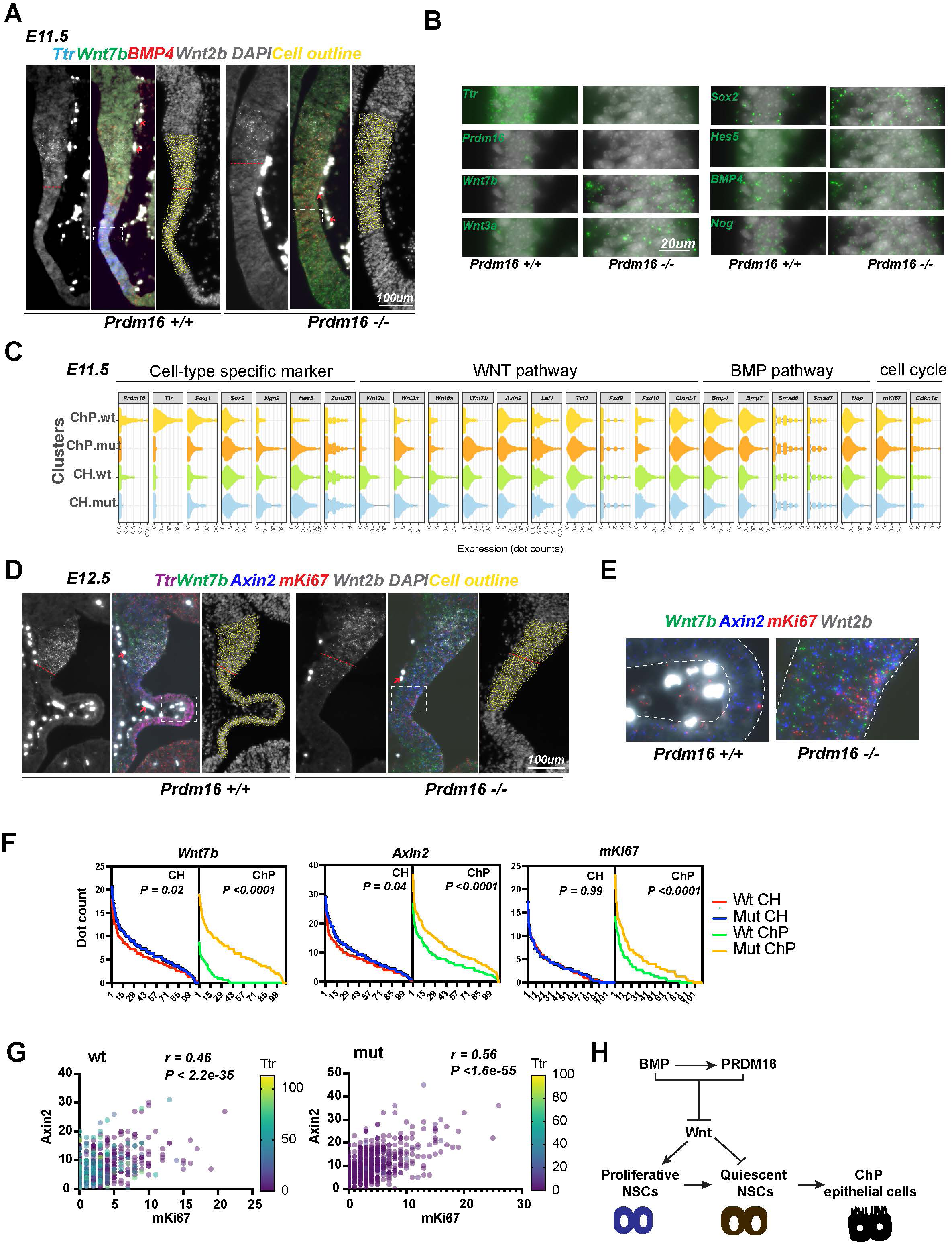
PRDM16 suppresses Wnt signaling and cell proliferation in ChP epithelial cells. (**A**) SCRINSHOT images of E11.5 wild-type and *Prdm16* mutant brains probed for *BMP4*, *Ttr*, *Wnt7b* and *Wnt2b*. Red arrows indicate autofluorescent blood cells (non-specific signal in all channels). Cells quantified are circled in yellow, and the red dashed line marks the CH and ChP boundary. (**B**) Enlarged views of the boxed region in (A), showing increased expression of Wnt and neural genes and decreased *Ttr* and *Prdm16* in mutants. (**C**) Violin plots displaying dot counts for the 24 genes in 330 CH and 330 ChP cells from three wild-type and three *Prdm16* mutant samples. (**D**) SCRINSHOT images of E12.5 wild-type and *Prdm16* mutant brains with probes for *Axin2*, *Ttr*, *Wnt7b*, *Wnt2b* and *mKi67*. (**E**) Enlarged views of the boxed areas in (D), with ChP regions outlined by dashed lines. (**F**) Distribution of dot counts for indicated genes in wild-type (*wt*) and *Prdm16* mutant (*mut*) CH and ChP cells. X-axis: 110 cells per replicate; Y-axis: mean dot count across three biological replicates. P values calculated using area under the ROC curve (Prism Graphpad). (**G**) Multi-variant linear regression of *mKi67*, *Axin2* and *Ttr* expression across 660 CH and ChP cells per genotype. Pearson correlation and significance were calculated using Prism Graphpad. (**H**) Model illustrating the regulatory circuit through which PRDM16 promotes the transition from proliferative to quiescent NSCs, a prerequisite step for ChP epithelial cell specification.

Next, we quantified expression changes of these genes in *Prdm16* mutant CH and ChP cells. As expression of *Wnt2b* was relatively unchanged in the mutant (**Supplementary Fig. 1** and **Fig. 7A**), we used *Wnt2b*-expressing cells to define the CH cells and those expressing *Ttr* to define ChP cells in the wild-type brain slice. There were about 110 cells in each region of one brain slice. Because the mutant brain slice showed little *Ttr* signal, we only used the *Wnt2b*-expressing cells to define the anterior border between CH and neocortex and included 110 cells under the border for the CH cells, and the further 110 cells for the mutant ChP cells, as shown in the images with DAPI and cell outlines (**Fig. 7A and Fig. 7D**). We then measured the dot count of each gene in these 220 cells in each sample, and summarized the changes from three pairs of animals in the violin plots (**Fig. 7C** and **Supplementary Fig. 6A**). Expression of *Ttr* and *Foxj1* is severely reduced, while *Sox2*, *Hes5* and *Ngn2* all become upregulated in the mutant ChP, indicating a fate transformation of ChP epithelial to neural cells. This agrees with our immunostaining and conventional *in situ* results (**Fig. 1** and **Supplementary Fig. 1**). Similar to BMP4, other BMP components we tested including *BMP7*, *Smad6*, *Smad7 and Nog* are unchanged, suggesting that PRDM16 is not an upstream regulator of the BMP pathway.

Moreover, *Wnt3a* and *Wnt7b* are significantly upregulated in the *Prdm16* mutant ChP (**Fig. 7B-E** and **Supplementary Fig. 6A-C**). In line with this, the Wnt target gene *Axin2* is also upregulated, indicating aberrantly elevated Wnt signaling in the *Prdm16* mutant ChP. Thus, the normal function of *Prdm16* represses WNT signaling in the developing ChP.

### Levels of Wnt activity correlate with cell proliferation in the developing ChP and CH

We further assessed the relationship between Wnt signaling and cell proliferation in the ChP and CH at E12.5, by correlating expression levels of *mKi67* with *Wnt3a*, *Wnt7b* and *Axin2* levels in wild-type and mutant samples. A significant increase of *mKi67* expression in *Prdm16* mutant ChP cells at E12.5 is accompanied by significantly increased Wnt gene expression (**Fig. 7F** and **Supplementary Fig 6B-C**). In contrast, little or no increase of *mKi67* signal is found in the mutant CH, suggesting that the effect of PRDM16 is mainly cell-autonomous. We then performed linear regression and Pearson correlation analyses to assess correlations between ChP markers, BMP and Wnt genes. Interestingly, the level of *Axin2* best correlates the level of m*Ki67*, even better than that with other Wnt genes in both wild-type (r = 0.46; p < 2.2 E-35) and mutant (r= 0.56; p < 1.6 E-55) CH and ChP cells **(Fig. 7G** and **Supplementary Fig. 6D**). This result suggests that Wnt activity may be responsible for cell proliferation in the CH and ChP region. Supporting this finding, it was shown that ectopic Wnt signaling converts ChP cells into proliferative CH neural cells ^12^. Similarly, in the 4^th^ ventricle, two Wnt genes, *Wnt1* and *Wnt3a*, and *mKi67* all became ectopically expressed in *Prdm16* mutant ChP cells (**Supplementary Fig. 7**). Thus, PRDM16 also suppresses Wnt signaling in the hindbrain ChP.

Additionally, six cell-proliferation-related genes (Spc24, Spc25, Nuf2, Ndc80, Id3 and Mybl2) exhibited upregulation in *Prdm16* mutant ChP (**Supplementary Fig. 8**). Interestingly, all four NDC80 complex genes became upregulated, pointing to its potential role in promoting NSC proliferation. Taken together, our results suggest that specification of the ChP epithelium requires a process transitioning proliferating NSCs into quiescence, and that this process is mediated by a suppressive role of PRDM16 on Wnt signaling and cell cycle regulators.

## Discussion

In this study we reveal a novel molecular mechanism by which BMP signaling regulates cell proliferation and gene expression. We find that PRDM16 acts as a tethering factor to localize the SMAD4/pSMAD1/5/8 complex at specific genomic sites and facilitate the repressive role of the SMAD proteins. The genes co-repressed by PRDM16 and BMP signaling include those encoding Wnt pathway ligands and other cell proliferation regulators.

Combinatory activities of morphogens are exploited throughout animal development and tissue homeostasis. The crosstalk between BMP and WNT signaling is complex as it can be synergistic or antagonistic ^46^. Surprisingly, both types of effects exist in the specification of ChP epithelium. Here BMP signaling induces NSC quiescence, as evidenced by the presence of ectopic proliferating cells at the ChP in *Bmpr1a* mutant mice ^10^, while Wnt signaling promotes proliferation since the gain-of-function condition of beta-Catenin phenocopies *Bmpr1a* mutant animals ^12^. On the other hand, adding a low dose of Wnt activator to cell culture medium enhances programing efficiency of ChP epithelial cells induced by BMP4 ^47^, and loss-of-function of beta-Catenin resulted in under-developed ChP structure ^12^, suggesting that BMP and WNT signaling collaborate to promote ChP epithelial cell differentiation. Our work demonstrated that PRDM16 ensures the right balance of BMP and Wnt activity by maintaining a low level of Wnt gene expression in the developing ChP.

PRDM16 was shown to physically interact with SMAD3 ^23^ and antagonize TGF-β-induced cell cycle arrest ^48^. However, we did not detect stable PRDM16/SMAD4/pSMAD1/5/8 or PRDM16/SMAD4/SMAD3 complexes in co-immunoprecipitation assays (data not shown). Instead, sequential ChIP revealed that PRDM16 and SMAD4 bind to the same DNA molecule, supporting their cooperative function at co-occupied regulatory elements. Notably, SMADs when associated with PRDM16 compared to those without PRDM16, with SMADs preferentially bind palindromic motifs in the presence of PRDM16 and GC-rich SBE motifs in its absence, suggesting that PRDM16 modulates SMAD DNA-binding specificity and shifts SMAD regulatory output.

A recent study similarly reported that PRDM16 acts as a SMAD4 co-repressor during pancreatic ductal adenocarcinoma progression ^49^. There, SMAD4 stably binds the PRDM16 promoter and switches its co-factor from the co-repressor PRDM16 (under low TGF-β) to the co-activator SMAD3 (under high TGF-β), depending on TGF-β levels. This mechanism differs from our model. Our data indicate that high BMP4 levels promote SMAD4 and pSMAD1/5/8 binding to target loci, including *Prdm16*, resulting in both gene repression and activation. Notably, PRDM16 contributes to the repression function of the SMAD complex even under elevated BMP conditions.

Although PRDM16 has been shown to promote proliferation by antagonizing anti-proliferation activity of TGF-β and SMAD3 ^48^, our data demonstrate that PRDM16 collaborates with SMAD4 and pSMAD1/5/8 to repress proliferation genes, which is consistent with BMP-induced quiescence. Similar roles have been observed in other contexts, such as such as osteoblast differentiation and maturation ^50^. Supporting this, genomic profiling revealed that under high BMP4 and PRDM16 conditions, pSMAD1/5/8 and SMAD4 occupy numerous loci without SMAD3, suggesting PRDM16 primarily modulates BMP target genes in these settings.

Moreover, while it was reported that PRDM16 enhances Wnt signaling by stabilizing nuclear beta-Catenin in the craniofacial tissue ^26^, we detected the opposite outcome in the ChP and cultured NSCs, ectopic WNT gene expression in the absence of *Prdm16*, suggesting that PRDM16 represses Wnt genes in these contexts. Notably, PRDM16, Wnt and BMP co-exist in various developmental settings, such as craniofacial development ^21,26,51^, heart formation ^27,52,53^, limb patterning ^21,54,55^, adult intestinal stem cells ^56–58^ *etc*. We speculate a similar regulatory circuit is used in some, if not all, of these settings.

ChP epithelial cells derive from proliferating NSCs, without intermediate fate-commitment steps. This property endows ChP cells with higher plasticity and makes them more vulnerable to abnormal genetic and cellular change, to which high occurrence frequency of ChP tumors in fetuses and young children may be attributed ^59^. As an essential brain structure, the ChP releases cerebrospinal fluid (CSF) and acts as a brain-blood barrier ^60–62^. Its dysfunction has been linked with several types of human diseases including hydrocephalus, Alzheimer’s disease, multiple sclerosis ^62^. Deepening our knowledge on the development of ChP will provide new insights into potential therapeutics to prevent or treat ChP tumors and other ChP-related diseases.

**Footnote**: a study on the antagonism between PRDM16 and SMAD4 was published on Feb 24, 2023 (Hurwitz, 2023) after we posted our manuscript at BioRxiv and during our submission process.

## Materials and methods

### Animals

All animal procedures were approved by Swedish agriculture board (Jordbruks Verket) with document number Dnr 11553-2017 and 11766-2022. The *Prdm16*^cGT^ mice ^22^ were maintained by outcrossing with the FVB/NJ line.

### In situ hybridization

To make probes for conventional RNA in situ hybridization, genomic regions covering one exon or full-length cDNA were PCR amplified to generate fragments with restriction enzyme overhangs. The sequences of all oligos were included in Supplementary table 1. The fragments were inserted to the pBluescript SK(II) vector. In vitro transcription was performed as previously described (He et al., 2021). The mouse brains at defined ages were dissected and fixed for 12 hours in 4% PFA, dehydrated in 25% sucrose, cryoprotected and embedded in O.C.T. The brain samples were then sectioned at 18 µm thickness on Leica cryostats CM3050s. RNA *in situ* hybridization was performed using digoxigenin-labeled riboprobes as described previously. Detailed protocols are available upon request. Images were taken using a Leica DMLB microscope.

### Immunostaining

Immunostaining was performed according to standard protocols as previously used ^63^. For EdU and BrdU labeling, EdU (5-ethynyl-2′-deoxyuridine) and BrdU (5-bromo-2’-deoxyuridine) (5-20 µg/g of body weight) were injected into the peritoneal cavity of pregnant mice at desired ages. EdU incorporation was detected with the Click-iT assay (Invitrogen) according to the manufacturer’s instructions. BrdU incorporation was measured by immunostaining using an antibody against rat-BrdU (Abcam). Imaging was taken on a Zeiss confocal microscope. ZEN (ZeissLSM800), ImageJ (NIH) and Photoshop (Adobe) were used for analysis and quantification.

### NSC culture

Control and mutant embryonic cortices from E13.5 animals were dissected and dissociated into single cell suspension and digested with Accutase (Sigma). Cells were maintained in proliferation media (STEMCELL Technologies). Lentivirus expressing Flag-*Prdm16* and a puromycin resistant gene in the pCDH vector was produced in 293J4 cells and used to infect a wild-type NSC line derived from E13.5 forebrain, and the selected NSCs were maintained in puromycin-containing medium for three passages before puromycin withdrawal.

### RT-qPCR

*Prdm16_OE*, wild-type and mutant NSCs were cultured and treated with or without BMP4 (Sigma). After 48-hour treatment, total RNAs were extracted using TRIzol reagent (Invitrogen). Total RNA was further cleaned with Turbo DNase (Ambion) and used in reverse-transcription with RT master mix (ThermoFisher). To ensure the absence of genomic DNA, control qPCR was performed on the mock-reverse-transcribed RNA samples. The list of qPCR primers is included in supplementary table 1.

### SCRINSHOT

Brain sections were prepared in the same way as for regular *in situ* hybridization (described above). The SCRINSHOT experiments were carried out according to the published method ^45^. In brief, 3 padlock probes and 3 corresponding detection probes were designed for each gene of interest. The slides with cryo-sectioned brain slices were pretreated at 45°C to reduce moisture and fixed in 4% PFA in 1X PBS, followed by washing in PBS Tween-20 0.05% twice. Permeabilization of tissues were done by washing slides in 0.1M HCl for 2mins 15s, followed by washing in PBS Tween-20 0.05% twice. Then a stepwise dehydration was performed for the slides in 70%, 85% & 100% Ethanol and air. The SecureSeal hybridization chamber (GRACE BIO-LABS, 621501) was then mounted to cover each pair of control and *Prdm16* mutant brain slices. Samples were then blocked in a probe-free hybridization reaction mixture of 1XAmplifase buffer (Lucigen, A1905B), 0.05M KCl, 20% Formamide deionized (Millipore S4117), 0.1uM Oligo-dT, 0.1ug/ul BSA (New England Biolabs, B9000S), 1U/ul RiboLock (Thermo, EO0384), and 0.2ug/ul tRNAs (Ambion, AM7119). Then hybridization of padlock probes was done by incubating samples with padlock probes (with the concentration of each one 0.01uM) mixed in blocking reagents used before (no oligo dT used in this step). The slide was then put into a PCR machine to denature at 55°C for 15 minutes and hybridize at 45°C for 120 minutes. Padlock probes were then ligated by using SplintR ligase (NEB M0375) at 25°C for 16 hours followed by RCA (rolling cycle amplification) at 30°C for 16 hours by using phi29 polymerase (Lucigen, 30221–2) and RCA primer1. Then a fixation step was applied to stabilize

RCA product in 4% PFA for 15 minutes followed by washing in PBS Tween-20 0.05%. Then the hybridization of the first 3 genes was done by mixing all 3 3’ fluorophore-conjugated detection probes of each gene in reaction reagent (2XSSC, 20% Formamide deionized, 0.1ug/ul BSA, and 0.5 ng/µl DAPI (Biolegend, 422801) followed by hybridization at 30°C for 1 hour. The slides were washed in 20% formamide in 2X SSC and then in 6X SSC, followed by dehydration in 70%, 85%, 100% Ethanol until the chamber was removed. Then the samples were preserved in SlowFade Gold Antifade mountant (Thermo, S36936) and kept in dark before imaging. After image acquisition, the first 3 detection probes were removed by using Uracil-DNA Glycosylase (Thermo, EN0362), and the slides were ready for the next round of detection probe hybridization. The procedure was repeated until all genes were hybridized and imaged.

Images were acquired with Zeiss Axio Observer 7 fluorescent microscope with an automated stage setting to fix imaging region for different hybridization rounds. Image analysis was done according to the published method ^45^. In brief, the DAPI channel from each round was extracted to measure the shift of imaging. The images were then aligned for all gene channels in Zen by Creating Image Subset with the shifting value. After alignment, images were exported into TIFF files and the threshold analysis was carried out for individual channel one by one in CellProfiler with scripts provided by the published method (Alexandros et al. 2020). Then nuclear segmentation was done manually in Fiji ROI manager to obtain nuclear ROIs, followed by an expansion of 2 um in CellProfiler to obtain cell ROIs. Then signal dots were count in these cell ROIs for each gene in CellProfiler and Fiji. Summary of the dot counts for each gene was exported to Excel files.

### CUT&TAG

CUT&TAG was performed according to the published method ^64^. In brief, *Prdm16*_*OE* and *Prdm16* mutant NSCs were cultured in NeuroCult™ Proliferation Kit (Stem cell technologies, 05702) for 3 days with or without BMP4 at concentration of 25ng/ml medium (Sigma, H4916). Cells on day 3 were then shortly rinsed with 1X PBS and resuspended in 1ml cold NE1 buffer on ice. Nuclei were lightly fixed with 0.1% formaldehyde to 1ml PBS at RT for 2 minutes and neutralized with 75mM Glycine. Nuclei were then washed for 3 times, resuspended in 1mL wash buffer and mixed with 90uL Concanavalin A-coated magnetic beads (Polyscience 86057). The nuclei/beads mix was then blocked with 800 uL cold Antibody buffer for 5 minutes and resuspended in 1.2 mL Antibody buffer and aliquoted into 8 tubes with 150 uL each. 2 uL Anti-PRDM16 (Generous gift from Bryan Bjork lab) or IgG (SIGMA-ALDRICH, I5006) was added into each tube for an overnight incubation at 4°C. The beads/nuclei/antibody mix was then washed with Dig-wash buffer and incubated with secondary antibody (1:100 dilution) for 1 hour. After further washes with the Dig-wash buffer. a pA-Tn5 adaptor complex in Dig-300 buffer was added to the beads for 1 hour reaction. The beads were then washed with Dig-300 buffer before the incubation with 300ul Tagmentation buffer at 37°C for 1 hour. Then 10ul 0.5M EDTA, 3uL 10% SDS and 2.5uL 20 mg/mL proteinase K (Invitrogen) were used to stop the reaction. Then the resultant DNA was purified with DNA clean & Concentrator kit (ZYMO Research D4013) and eluted in 25uL Elution buffer. To generate libraries, 21uL fragment DNA was mixed with 2uL 10uM Universal i5 primer, 2uL 10uM uniquely barcoded i7 primer and 25uL PCR master mix (NEB Phusion® High-Fidelity PCR Master Mix with HF Buffer, M0531). The PCR condition was as follows: 72°C 5min, 98°C 30s, repeat 12 times (98°C 10s, 63°C 10s), 72°C 1 min and hold 4°C. The libraries were cleaned with standard Ampure XP beads as previously described. Libraries from four biological replicates were produced for each condition.

### ChIP

ChIP was performed as previously described ^65^. For each ChIP reaction, 10 million *Prdm16_OE*, Prdm16_KO NSCs with or without 3-day BMP4 treatment were fixed, lysed, sonicated and made into chromatin extract. After precleared with gamma-bind-G beads, the chromatin extract was incubated with 2ug PRDM16, 2ug SMAD4 (Proteintech, 10231-1-AP), 2ug pSMAD1/5/8 (Millipore, AB3848-1) or 2ug SMAD3 (abcam, ab227223) in each ChIP reaction. In sequential ChIP assays, the same chromatin lysate was precleared and used, but with more antibodies in each reaction: 5 ug of IgG or PRDM16 antibody in the first round of immunoprecipitation. The elutes were then precleared again using gamma-bind G beads, divided into two equal halves and immunoprecipitated with 2 ug of IgG or the SMAD4 antibody. In ChIP-qPCR experiments, 2ug SMAD4 and 2 ug c-FOS antibodies (Invitrogen, MA5-15055) were used in each reaction. The precipitated DNA was reverse cross-linked, and then purified using the Qiagen PCR purification kit.

### Computation analyses

#### ChIP-seq libraries and analyses

0.2% input and ChIPed DNA were made into libraries using the NEBNext Ultra™ II DNA Library Prep Kit and sequenced on the Illumina Nextseq500 platform. Three replicates of ChIP-seq samples, after the adaptor trimming by Trimmomatic, were mapped to the UCSC *Mus musculus* (mm10) genome assembly using Bowtie2 with the default parameters. The uniquely mapped reads (with mapping quality >= 20) were used for further analyses. The peaks were called by HOMER (v4.10) ^66^. The reproducibility between replicates was estimated by Irreproducibility Discovery Rate (IDR), using the HOMER IDR pipeline (https://github.com/karmel/homer-idr). As suggested by the Encode IDR guideline, we used a relatively relaxed parameter “-F 2 -fdr 0.3 -P .1 -L 3 -LP .1” for the true/pseudo/pooled replicates by the HOMER peak calling. The final confident peaks were determined by an IDR < 5%. The peaks that were overlapped with mm10 blacklist were also removed.

#### CUT&TAG analyses

The CUT&TAG samples of *Prdm16_OE* and *Prdm16* mutant NSCs in four replicates, and of Prdm16 from CH and ChP area at stage 12.5 embryos in three replicates, were mapped to mm10 genome assembly using Bowtie2 (bowtie2 --end-to-end --very-sensitive --no-mixed --no-discordant --phred33 -I 10 -X 700). The CUT&TAG coverage for these samples were generated by bedtools genomeCoverageBed and normalized to the library depths to give a read per million per base (RPM). By using the peak caller SEACR ^33^, which was designed for calling peaks from sparse chromatin profiling data such as CUT&TAG [22], we performed peak calling with FDR < 10% for each replicated sample (SEACR_1.3.sh normalized_coverage 0.1 non stringent output_peaks). To identify confident peaks, we selected the common peaks, which were called from all replicates for each condition. Peak overlapping analysis was performed by Homer mergePeaks function with default parameters. Peak overlap analysis utilized HOMER’s mergePeaks function, specifically employing the default ’-d given’ parameter. This approach ensures that only peaks directly overlapping with each other are merged, maintaining the precise distances between peaks as provided in the input peak calls. This method is particularly beneficial for retaining the integrity of the original peak boundaries, avoiding any alterations to their relative positions. *De novo* motif discovery from SMAD4 and pSMAD1/5/8 peaks in *Prdm16_OE* NSCs and Prdm16 mutant NSCs were performed by MEME-ChIP software ^67^ (parameters: -ccut 100 -meme-p 5 -dna -meme-mod anr -minw 5 - maxw 15 -filter-thresh 0.05).

The CUT&TAG samples of H3K4me3 BMP4-treated and non-treated *Prdm16*_*OE* and *Prdm16*_KO cells in 2 or 3 replicates were mapped to mm10 genome assembly using Bowtie2 (bowtie2 --end-to-end --very-sensitive --no-mixed --no-discordant --phred33 -I 10 -X 700). Differential analysis of TSS up-stream and down-stream 500 bp between *Prdm16_OE* BMP-treated vs non-BMP-treated and Prdm16_KO vs *Prdm16_OE* BMP-treated were performed by Limma R package ^68^. Gene Ontology (GO) enrichment analysis of up-regulated and down-regulated regions was performed by PANTHER ^69^.

#### scRNA-seq analyses

The scRNA-seq data of the developing mouse brain were obtained from^41^. The counts per cell were normalized by logNormCounts function in Bioconductor package “scater” and the normalized expression data per cell were used to generate gene expression violin plots. To test whether Prdm16 target gene sets of Prdm16 with and without BMP4 and ChP E12.5 are significantly enriched among the scRNA-seq gene mean expression in each identified cell type cluster, the gene set enrichment analysis (GESA) “fgesa” Bioconductor package was used [ref: Korotkevich G. et al. Fast gene set enrichment analysis. bioRxiv, 2021. http://biorxiv.org/content/early/2016/06/20/060012].

## Supporting information

Reagent list

## Acknowledgements

We thank the animal experimental core facility and the imaging facility of Stockholm University and Bioinformatics and Expression analysis core facility at Karolinska Institute, Sweden, for their service and support. We thank Christos Samakovlis and his lab members for technical help on SCRINSHOT experiments. We also appreciate technical help from Adrian Martinez Martin. J.W. is funded by Australian Research Council Centre of Excellence for the Mathematical Analysis of Cellular Systems (CE230100001) and ANU Future Scheme. The project was supported by the research project grant from Swedish Research Council (Vetenskapsrådet, 2020-03543) and the research grants from Swedish Cancerfonden (CAN 2017/529 and 20 1046 PjF 01 H) to Q.D.

## Author contributions

Q.D. conceived and designed the project. L.H. performed all of the experiments. J.W performed all computational analysis. Q.D., J.W. and L.H. analysed and interpreted the data. Q.D. wrote the manuscript with input from L.H. and J.W.

## Data availability

The CUT&TAG and ChIP-seq data produced in this study have been deposited to Gene Expression Omnibus (GEO accession number: GSE275758).

## Figure legend

**Supplementary Figure 1.**
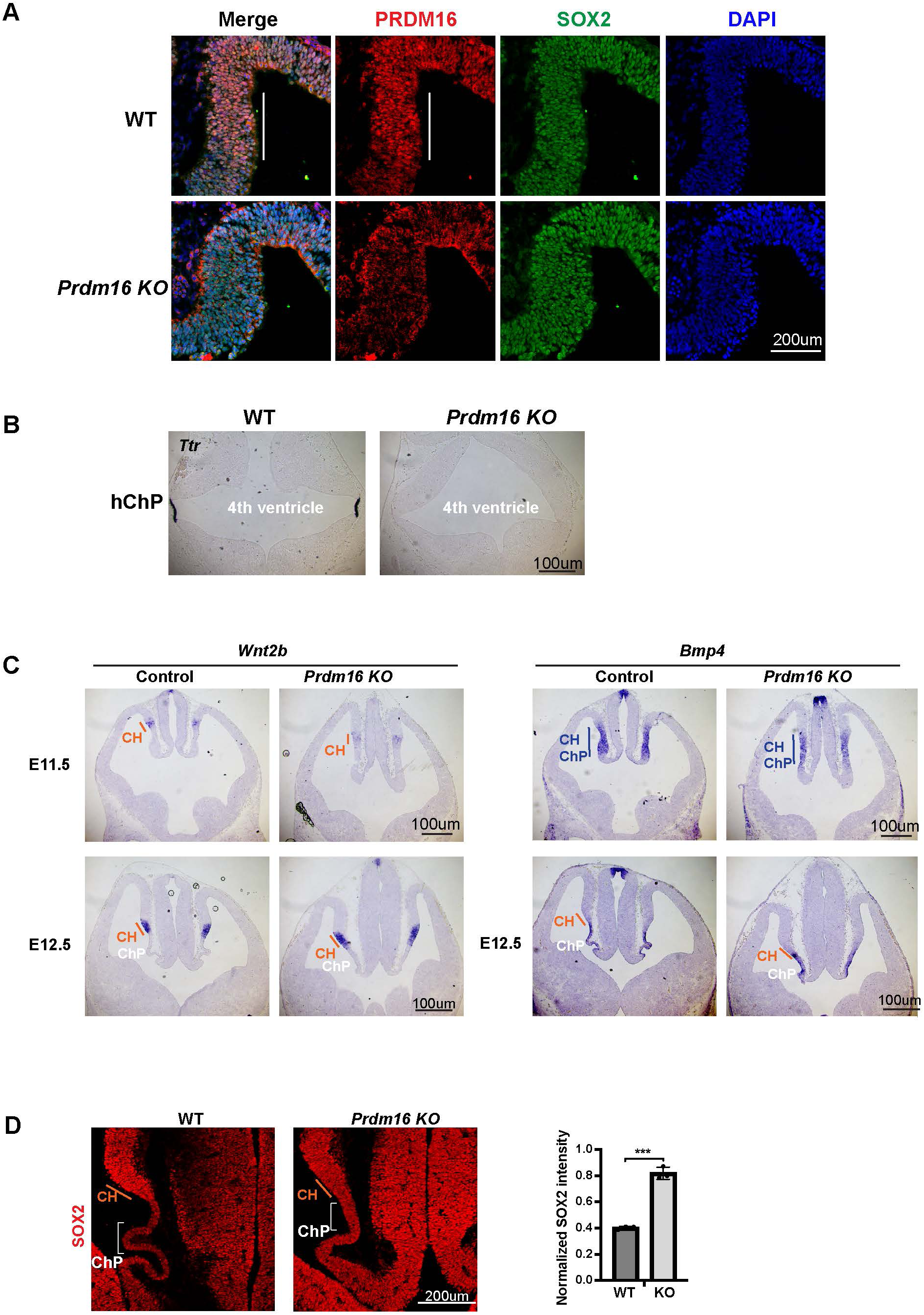
*Prdm16* depletion affects ChP epithelial specification without altering dorsal midline patterning. (**A**) Immunostaining of PRDM16 and SOX2 on E10.5 control and homozygous brain sections. ChP regions are outlined in white. (**B**) RNA in situ hybridization for *Ttr* in E12.5 control and Prdm16 null mutant hindbrains. (**C**) RNA in situ hybridization using *Wnt2b* and *Bmp4* probes on control and mutant forebrains at E11.5 and E12.5. (**D**) SOX2 immunostaining on E12.5 brain sections. ChP areas are marked with brackets. Relative SOX2 signal intensity between ChP and neurogenic regions is quantified. Error bars represent standard deviation (SD), n = 3. Statistical analysis by unpaired t-test (***P < 0.001; **P < 0.01; *P < 0.05; n.s., not significant).

**Supplementary Figure 2.**
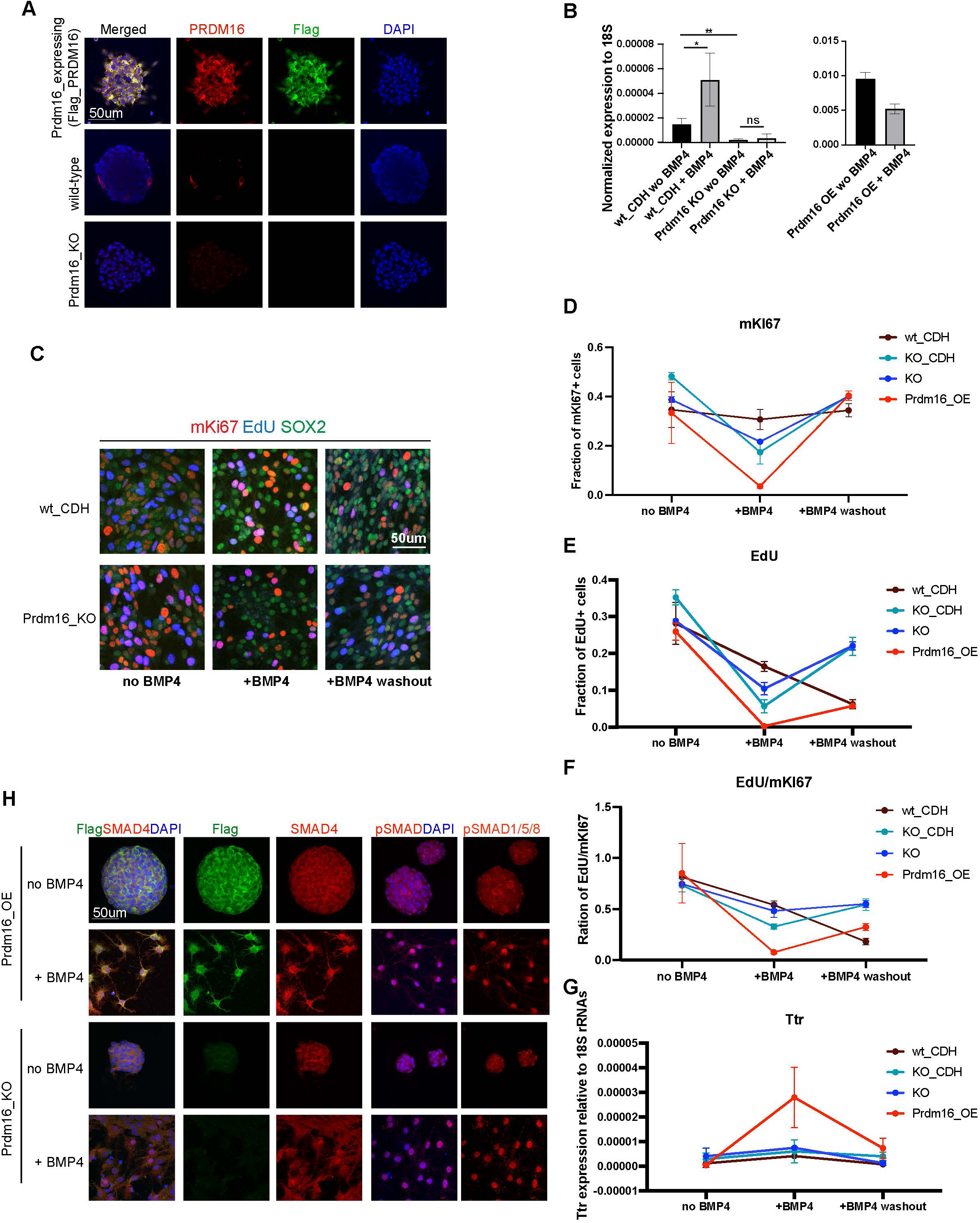
NSC culture assay reveals PRDM16 function in NSCs. (**A**) Immunostaining for PRDM16 and FLAG with DAPI in NSCs expressing *Flag_Prdm16*, wild-type (WT) NSCs, and *Prdm16_KO* NSCs. PRDM16 is sparsely expressed and cytoplasmic in WT NSCs, whereas FLAG_PRDM16 is robustly expressed in both cytoplasm and nucleus. (**B**) RT-qPCR measurement of *Prdm16* mRNA levels normalized to the 18S rRNAs in the indicated genotypes and conditions, from three biological replicates, each with two technical replicates. Statistical significance was calculated using unpaired t-test (***, p<0.001; **, p< 0.01; *, p<0.05; n.s., non-significant). (**C**) Immunostaining of dissociated NSCs on Matrigel for mKI67, EdU and Sox2. WT_CDH denotes WT NSCs infected with the pCDH vector. (**D-F**). Quantification of marker-positive cell fractions under three culture conditions. KO_CDH indicates *Prdm16_KO* NSCs infected with the pCDH vector. (**G**) RT-qPCR measurement of *Ttr* mRNA levels normalized to 18S rRNAs, from three biological replicates. (**H**) Immunostaining for SMAD4, pSMAD1/5/8 and FLAG together with DAPI in NSCs. Nuclear localization of SMAD4 and pSMAD1/5/8 are enhanced following BMP4 treatment.

**Supplementary Figure 3.**
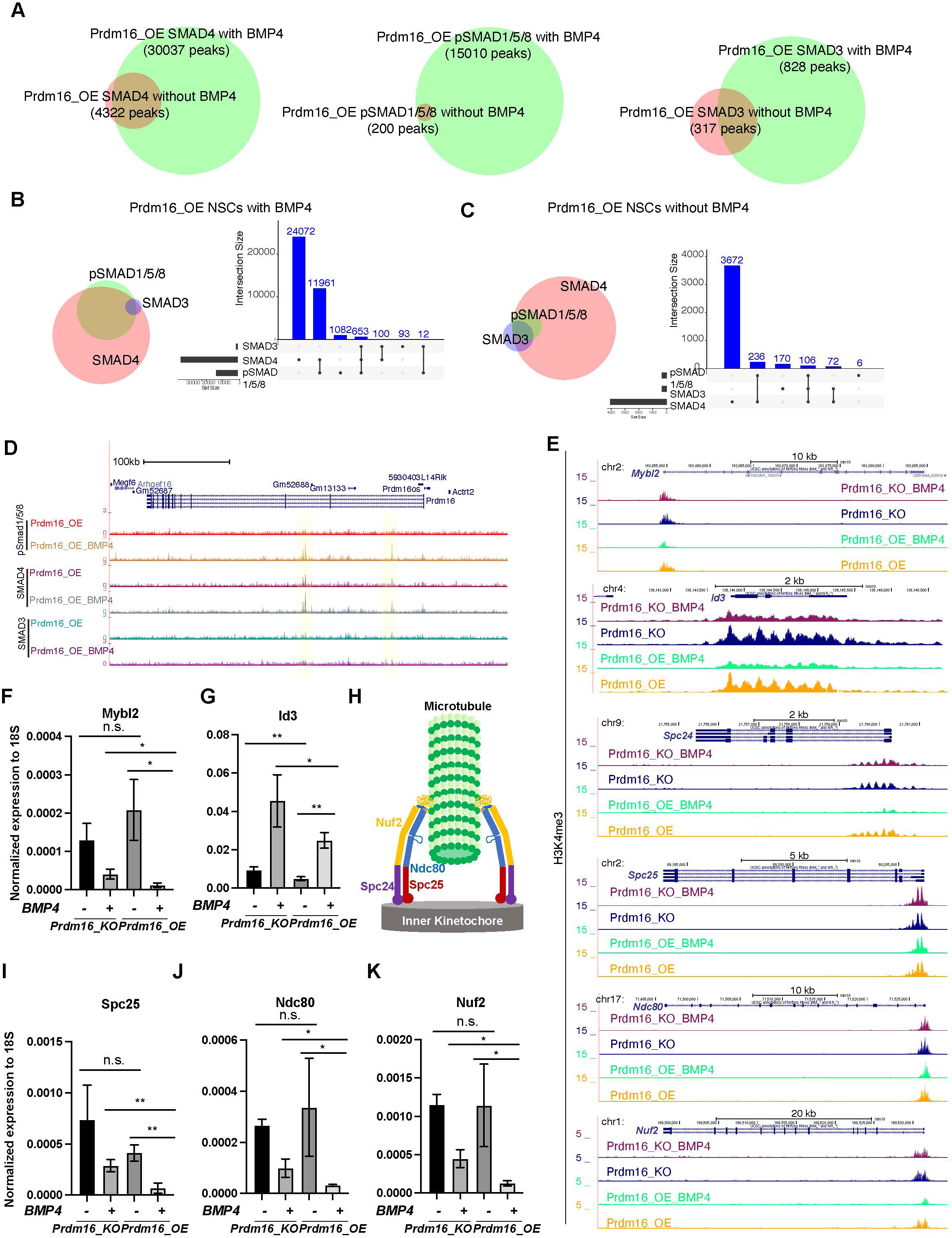
Overlapping analysis of SMAD4, pSMAD1/5/8 and SMAD3 CUT&TAG peaks. **(A)** Venn diagrams showing overlapping SMAD4, pSMAD1/5/8 and SMAD3 ChIP-seq peaks in *Prdm16_OE* NSCs with or without BMP4. (**B-C**) Overlap Venn diagrams and bar plots for SMAD4, pSMAD1/5/8 and SMAD3 ChIP-seq peaks in *Prdm16_OE* NSCs with BMP4 (B) and without BMP4 (C). (**D**) Genome browser tracks of the *Prdm16* locus showing increased SMAD4 and pSMAD1/5/8 binding at the two highlighted enhancer regions. (**E**) Genome browser tracks showing the H3K4me4 CUT&TAG signals at the indicated loci. **(F-G, I-K**). RT-qPCR measurement of the indicated genes normalized to 18S rRNAs, from three biological replicates. Error bars represent standard deviation (SD). (**H**) Schematic of the NDC80 complex. Statistical significance is calculated using unpaired t-test. ***, p<0.001; **, p< 001; *, p<0.05; n.s., non-significant.

**Supplementary Figure 4.**
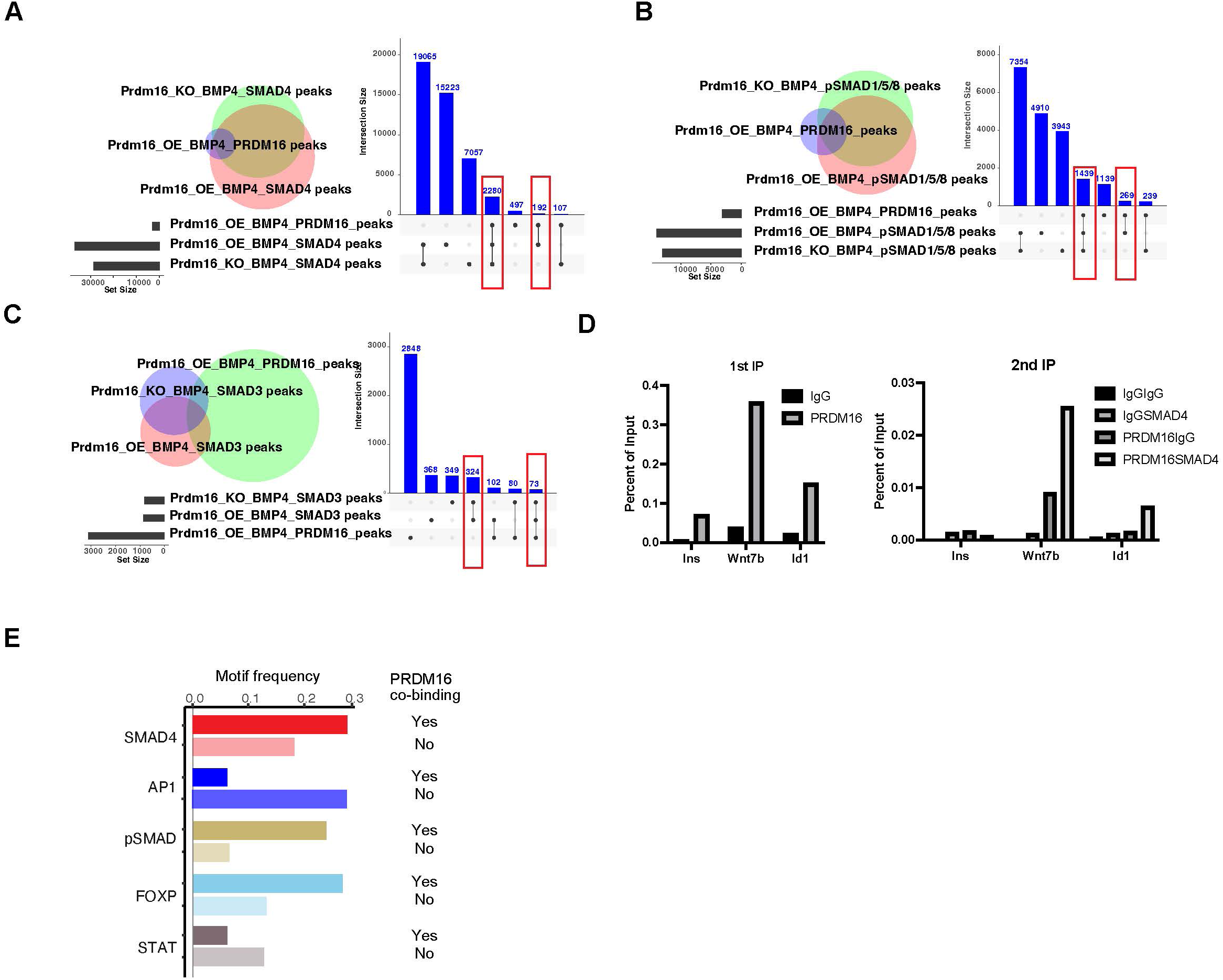
PRDM16 assists SMAD4/pSMAD1/5/8 genomic binding to their co-bound sites. (**A-C**) Venn diagrams and interaction plots showing overlap between PRDM16 CUT&TAG peaks in *Prdm16_OE* cells with BMP4, SMAD4 (A), pSMAD1/5/8 (B), SMAD3 (C) ChIP-seq peaks in *Prdm16_OE* and *Prdm16_KO* cells with BMP4. (**D**) Sequential ChIP-qPCR using IgG or PRDM16 antibodies followed by SMAD4 antibody. The *Ins* gene serves as a negative control. (**E**) Frequency of each DNA motif in SMAD4/pSMAD1/5/8 ChIP-seq peaks with or without PRDM16 co-binding in BMP4 treated *Prdm16_OE* cells.

**Supplementary Figure 5.**
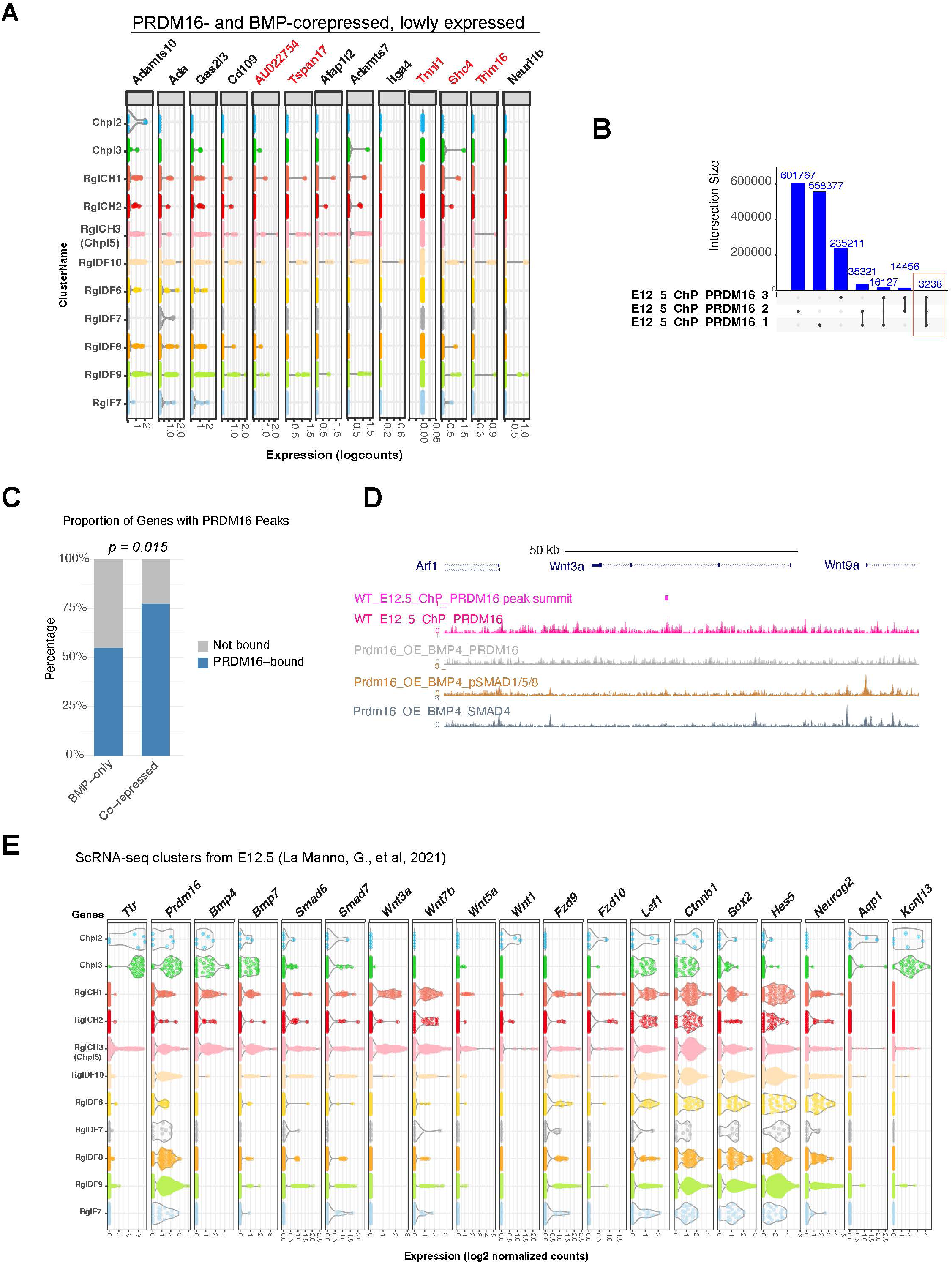
Wnt genes are potential targets regulated by PRDM16. **(A)** Violin plots of scRNA-seq expression at E12.5 for the indicated genes in cell type clusters (reanalyzed from La Manno et al., 2021 as in Figure 6A). Genes shown here have lower expression than those in main Figure 6A. Red text indicates genes without PRDM16 binding in E12.5 ChP. **(B)** Intersection plot of PRDM16 CUT&Tag peaks across three biological replicates; commonly shared peaks are highlighted. (**C**) Bar graph showing proportion of genes with PRDM16 binding in E12.5 ChP. (**D**) Genome browser view of Wnt3a locus showing PRDM16 CUT&Tag and SMAD ChIP-seq tracks. (**E**) Violin plots of gene expression across reanalyzed cell type clusters.

**Supplementary Figure 6.**
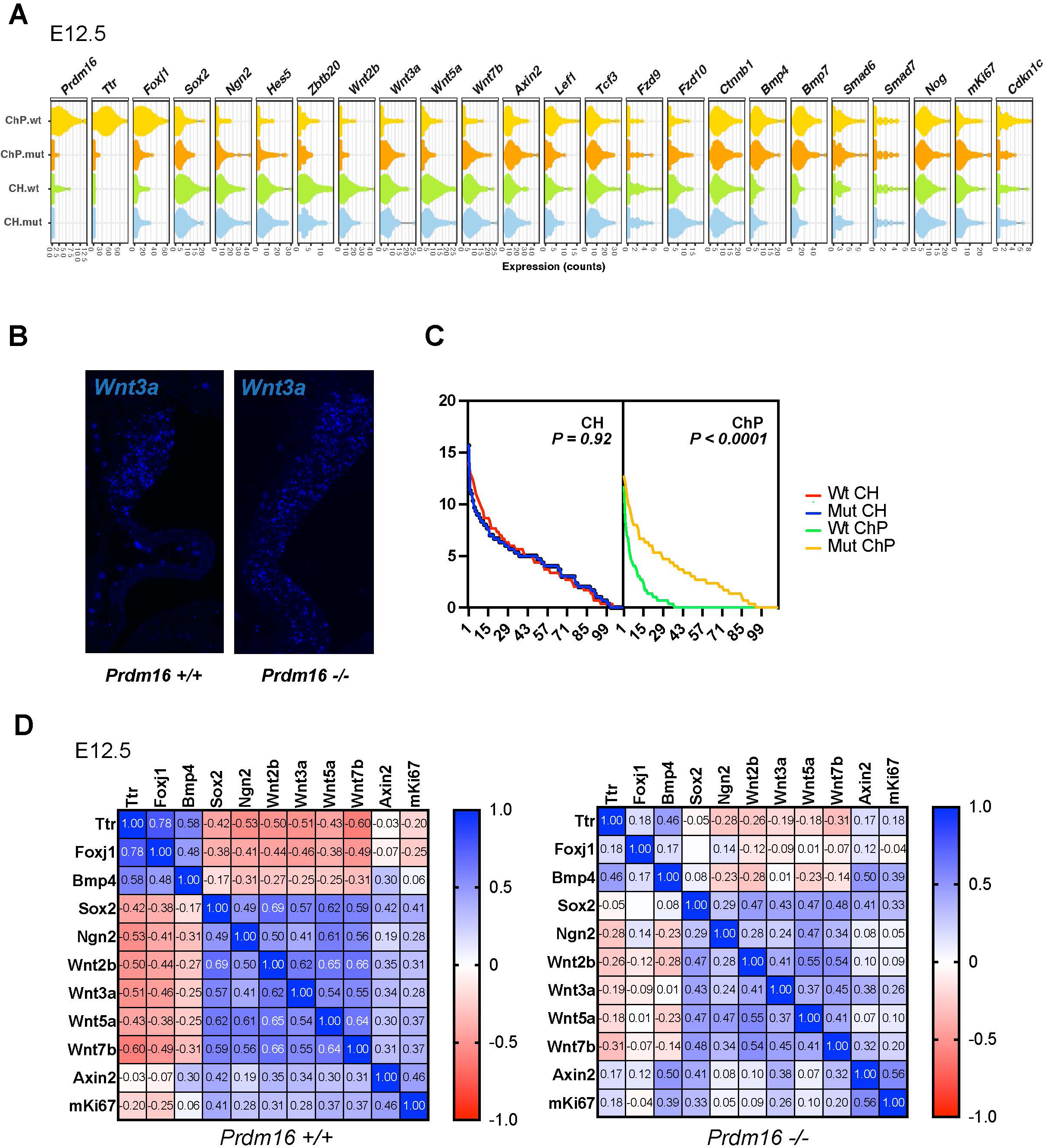
Altered expression of Wnt pathway and proliferation genes in *Prdm16* mutant ChP cells. (**A**) Violin plots showing expression changes of 24 examined genes (including ChP, neural, BMP pathway, Wnt pathway genes and *mKi67*) across 110 CH and 110 ChP cells per sample (330 cells per genotype, 3 WT and mutant pairs). (**B**) SCRINSHOT images showing *Wnt3a* mRNA in WT and *Prdm16* mutant brains. (**C**) Dot quantification of *Wnt3a* expression in CH and ChP regions across three replicates. X-axis: cell ID; Y-axis: average dot count. *P* value was calculated using area under the ROC curve in Prism Graphpad. (**D**) Heatmap of gene expression correlations. In WT cells, BMP genes cluster with *Ttr* and *Foxj1*, while Wnt genes cluster with *mKi67*. In mutants, BMP genes correlation with ChP markers is reduced, while Wnt-*mKi67* correlations is maintained.

**Supplementary Figure 7.**
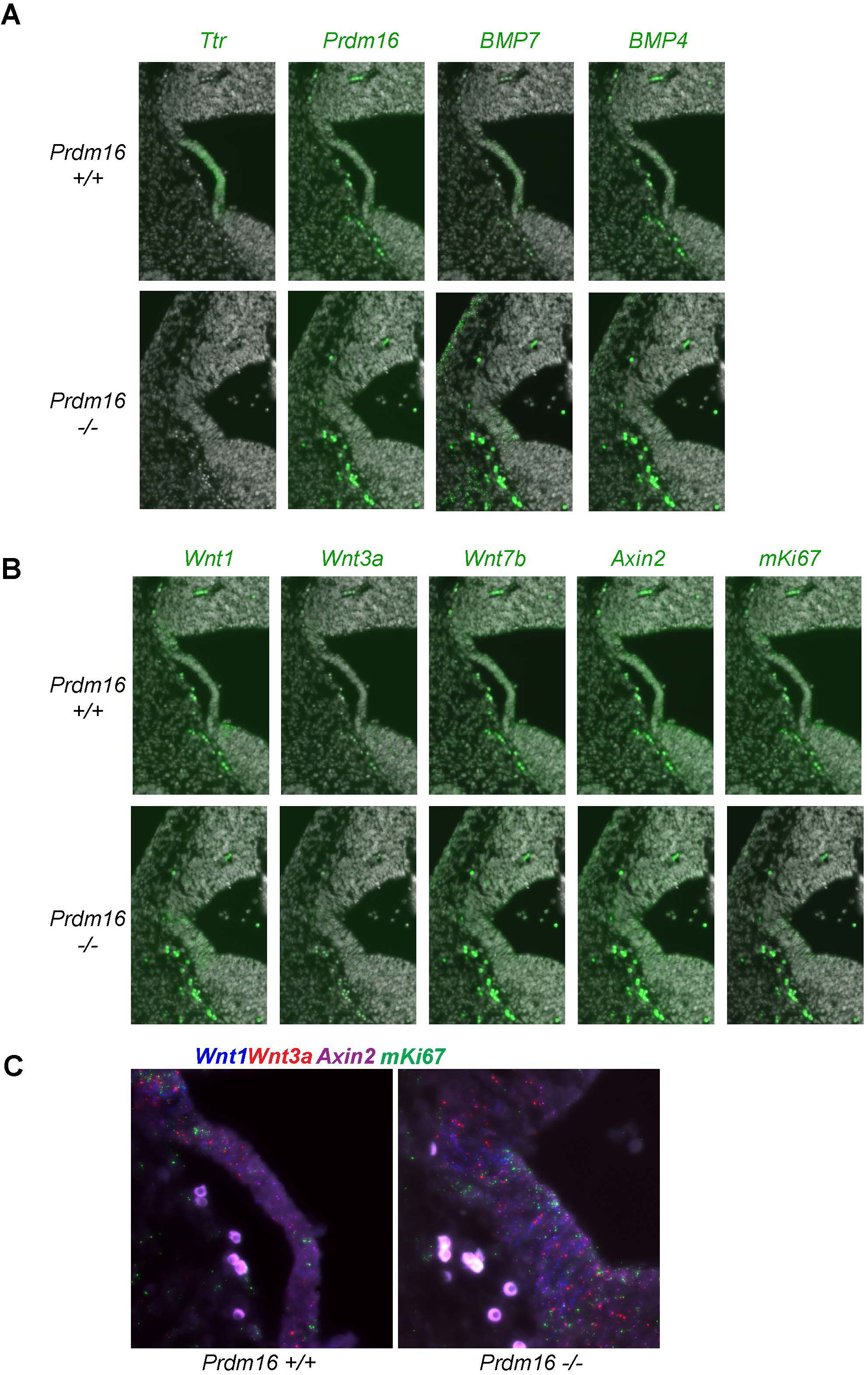
Expression of BMP and Wnt genes in the developing 4^th^ ventricle ChP. (**A-B**) SCRINSHOT images showing mRNA levels of ChP marker and BMP genes (A), and Wnt genes and *mKi67* (**B**), in wild-type and *Prdm16* mutant brains. Green: mRNA signals; grey: DAPI. **(C)** Enlarged view of SCRINSHOT images. Note the increased dot count for Wnt genes and *mKi67* in the mutant.

**Supplementary Figure 8.**
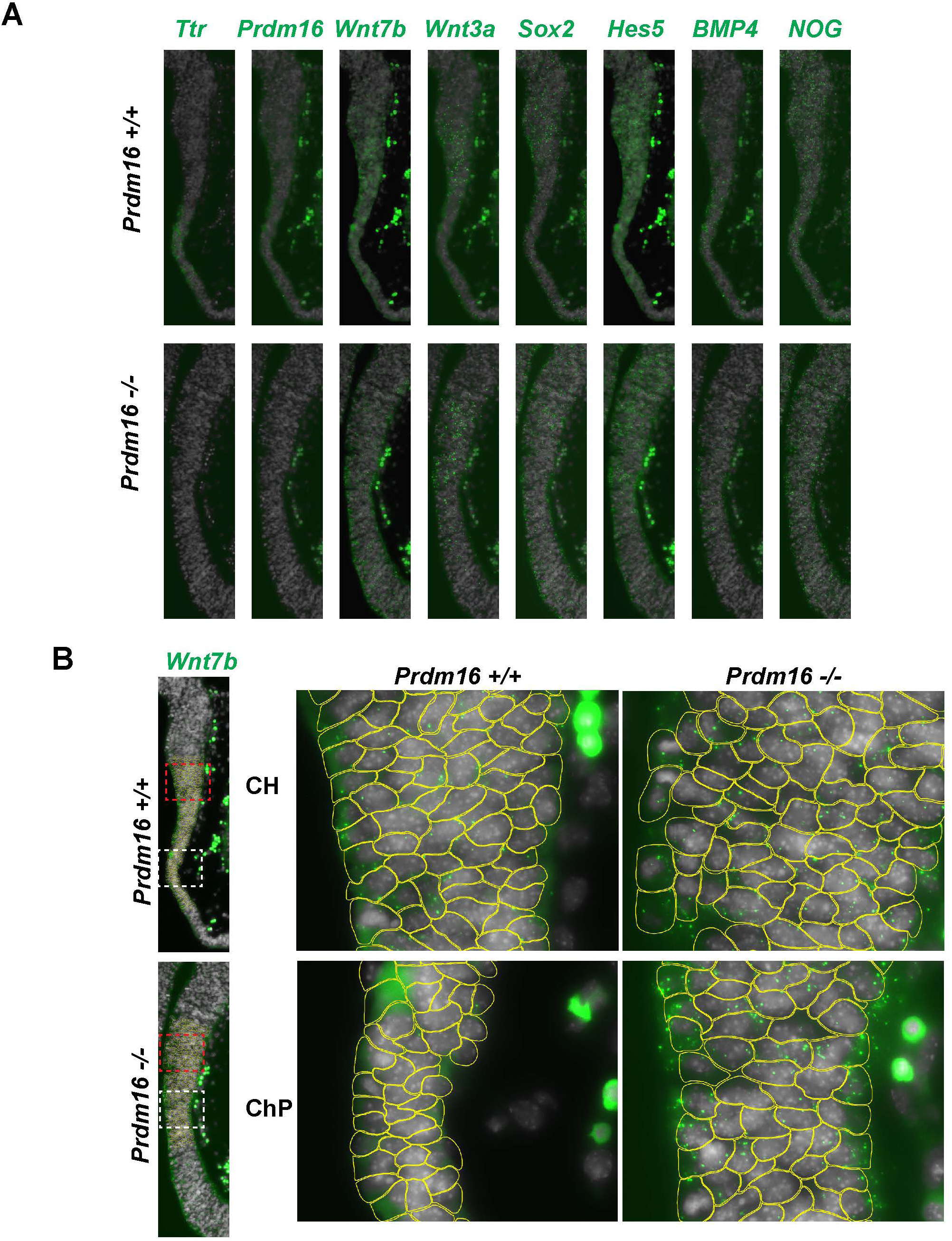
Increased expression of additional PRDM16 target genes in *Prdm16* mutant ChP. **(A)** Full images of SCRINSHOT signals for genes shown in Figure 7B. (**B**) Example of image segmentation for SCRINSHOT dot quantification.

**Supplementary Figure 9.**
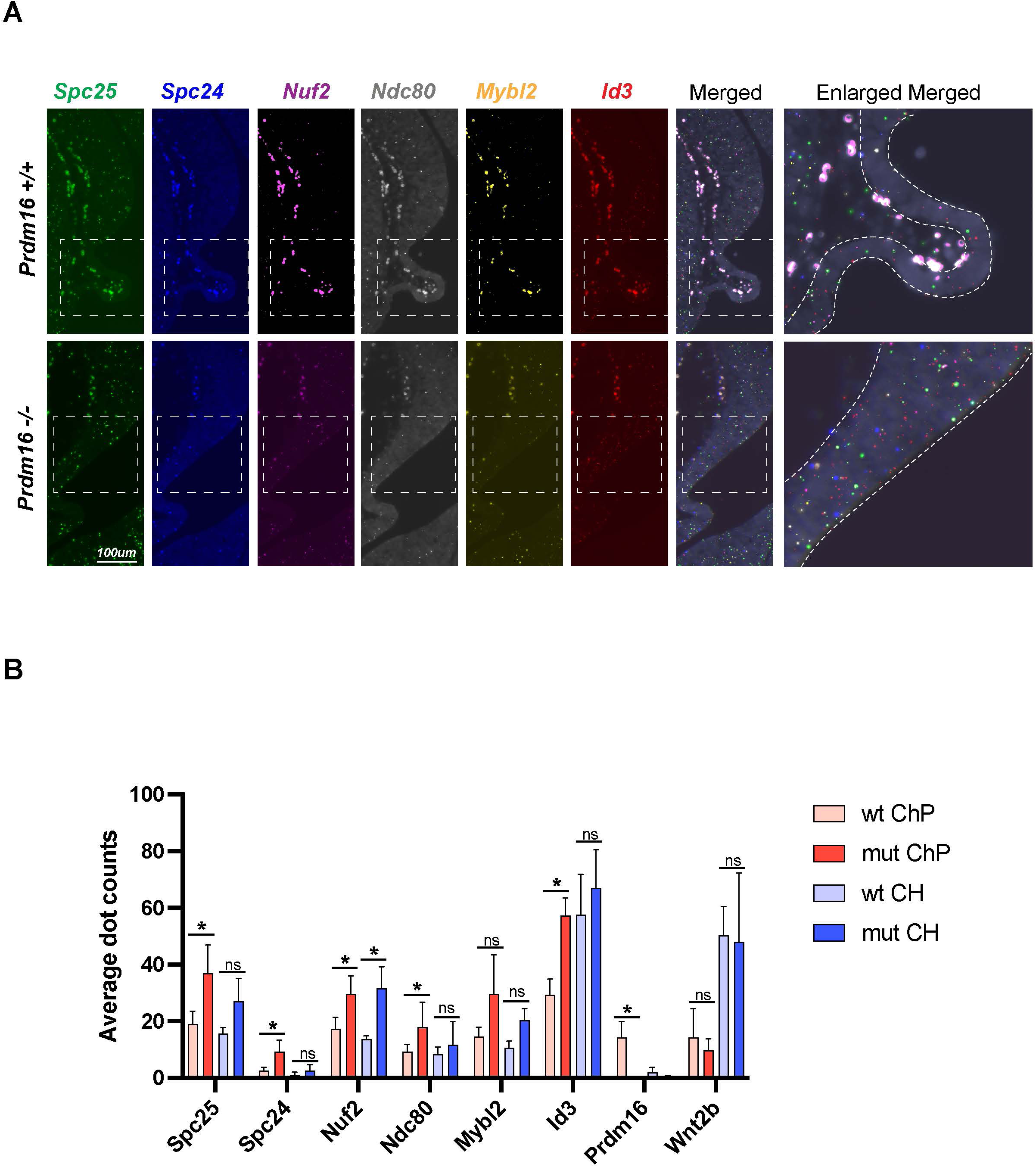
Increased expression of additional PRDM16 target genes in *Prdm16* mutant ChP. **(A)** SCRINSHOT images showing mRNA levels of indicated genes in wild-type and *Prdm16* mutant brains; enlarged views at right. Gene expression is elevated in mutants. (**B**) Quantification of dot counts in CH and ChP regions (110 cells each) across three replicates. *Prdm16* and *Wnt2b* are included as controls. Error bars denote SD. Statistical analysis via paired t-test (***P < 0.001; **P < 0.01; *P < 0.05; n.s., not significant).

